# Too hot or too disturbed? Temperatures more than hikers affect circadian activity of females in northern chamois

**DOI:** 10.1101/2023.02.03.527075

**Authors:** Lucie Thel, Mathieu Garel, Pascal Marchand, Gilles Bourgoin, Anne Loison

## Abstract

Recreational activities often result in a spatial and/or temporal activity shift in wildlife. With the concurrent development of outdoor activities and increase in temperatures due to climate change, mountain species face increasing pressures in terms of managing their activity pattern to limit both risk exposure and thermal discomfort. Using more than 15 years of long-term GPS and activity sensor data, we investigated how female northern chamois, *Rupicapra rupicapra rupicapra*, adjust their summer circadian activity to spatiotemporal variation in both temperatures and hikers’ presence. Chamois behaviour was more affected by high temperatures than by hikers’ presence. During the hottest days, they shifted their activity peak earlier in the morning, were more active at night, less active during daytime and had longer morning and evening peaks compared to the coldest days. Global activity was only slightly different during the hottest than the coldest days. Conversely, hikers’ disturbance had weak effects on activity levels and on the timing of activity peaks. This is especially true for temporal disturbance (weekdays versus weekends and public holidays), possibly because most weekdays in summer fell during school holidays. During the hottest conditions, the morning activity peak was shorter and the evening peak longer in females living in the most exposed areas compared to females living in the least exposed areas. One possible explanation for the overall low effect of hikers’ disturbance may be that behavioural changes buffering animals from high temperatures and hikers’ presence (e.g. moving away from trails) allow them to just marginally modify their activity pattern. In the context of ongoing socioenvironmental changes, it is critical to conserve habitats providing thermal refuges against summer heat and protection from disturbance to mitigate potential detrimental consequences.

## INTRODUCTION

A noticeable development and diversification of outdoor activities has been ongoing in the twenty-first century, especially in Western countries (Wheaton, 2010). While mountain ecosystems have been preserved until recently because of their low accessibility, they now face an increase in unaffiliated activities such as hiking, cross country skiing, trail running or climbing. These recreational activities are nonlethal, but humans can be consistently perceived as predators by wildlife (Frid & Dill, 2002), leading to a suite of negative physiological and behavioural responses (Arlettaz et al., 2007; Marchand et al., 2014; Larson, Reed, Merenlender, & Crooks, 2016). The consequences of repeated reactions to disturbance can be detrimental for several fitness-related parameters (Phillips & Alldredge, 2000; French, González-Suárez, Young, Durham, & Gerber, 2011). This is particularly true in mountain ecosystems, where the diversification of human activities challenges animals all year long, during both the period of energy acquisition from spring to autumn and the period of energy loss during winter (Lindstedt & Boyce, 1985). This increased pressure on the animals is accompanied by environmental modifications, including climate change, that are particularly strong in the Alps (Pepin et al., 2015). Temperatures here are rising rapidly in all seasons, with consequences such as earlier springs and plant cycles, more frequent summer heatwaves and droughts, and decreased duration of snow cover associated with an increased risk of frost for plants (e.g. Gobiet et al., 2014). Mountain animals are therefore concomitantly facing two major changes, namely an increased disturbance level and climate change.

To reduce their exposure to human disturbance, animals can adjust their behaviour in terms of movements, habitat use and selection, and time devoted to their different activities (foraging, resting, vigilance and social interactions). When facing a disturbance perceived as a risk, animals can modify their activities (e.g. from foraging to vigilance; Benoist, Garel, Cugnasse, & Blanchard, 2013; Xu, Gong, & Wang, 2021), and move away temporarily or even permanently (Rogala et al., 2011). They can also modify their activity pattern by delaying activities, such as foraging in disturbed places, from high-to low-risk periods (Bonnot et al., 2013). The benefits of such responses have been explained within the ‘risk allocation hypothesis’ framework (Lima & Bednekoff, 1999): the costs of lost opportunities to forage during risky periods could be reduced if, for instance, animals transfer some of their diurnal activity to night-time when human activities stop (e.g. Marchand et al., 2014; Bonnot et al., 2020). However, activity patterns are regulated, such as the critical foraging/rumination succession in ruminants (Hamel & Côté, 2008). Any forced modification of these patterns can negatively impact the fitness of individuals, for instance via an increased encounter risk with predators (Lesmerises et al., 2018) or mismatches in interactions with conspecifics (Greives et al., 2015),.

At the same time, animals need to deal with climatic constraints, and behavioural thermoregulation is among the most important mechanisms to avoid thermal stress. Indeed, temperatures influence space use (arctic ground squirrels, *Spermophilus parryii*, Long, Martin, & Barnes, 2005; in Mediterranean mouflon, *Ovis gmelini musimon* x *Ovis* sp., Marchand et al., 2015) and activity budgets (Japanese monkeys, *Macaca fuscata*, Watanuki & Nakayama, 1993), with animals having to trade between foraging and thermal cover to limit thermoregulatory costs while maintaining their energy balance (van Beest & Milner, 2013). In the same way as they do when avoiding human disturbance, animals can shift their activity peaks earlier in the morning and later in the evening, and increase their nocturnal activity during the hottest summer days (Bourgoin et al., 2008; Bourgoin et al., 2011). However, in temperate ecosystems, the night can be significantly shorter than the day during summer, and riskier when nocturnal predators are present. Thus, the transfer of the diurnal activity to night-time does not necessarily allow animals to fully compensate for the loss of foraging opportunities during the day and may, for instance, have consequences for mass gain (Garel, Loison, Gaillard, Cugnasse & Maillard, 2004) or predation risk (Gehr et al., 2017). These results highlight the importance of investigating full daily cycles to get a comprehensive picture of how animals may adjust their behaviour in the face of human-induced environmental changes (Marchand et al., 2014; Marchand et al., 2015).

Moreover, ongoing climate change and expansion of human activities in mountain ecosystems are also ultimately connected. An extended summer season or more frequent summer heatwaves in valleys may, for instance, increase the presence of humans in the mountains. Although predicting tourist numbers under scenarios of climate change is probably not straightforward (Beniston, 2003), this raises questions on animal responses to these interrelated human-induced environmental changes and on their potential cumulative or even multiplicative consequences for populations in terms of both immediate and compensatory responses (Schmeller et al., 2022). One critical challenge is therefore to understand whether the behavioural mechanisms used by animals to buffer against increasing temperatures may also provide an advantage against human presence or vice versa.

Here, we used GPS monitoring of 62 female northern chamois, *Rupicapra rupicapra rupicapra*, to investigate at a fine scale their behavioural responses to temporal and spatial stressors during summer, through the study of their activity pattern. Specifically, we assessed the day-to-day variation in temperatures and hikers’ presence for each chamois. We also computed an individual-specific index of spatial disturbance, combining data on the density of hiking trails in the summer home range of each chamois and the presence of hikers in each trail section. A diversity of activity metrics (e.g. daily activity levels, timing of activity peaks, length of activity peaks) allowed us to investigate the relative contribution of spatiotemporal variation in both temperatures and human disturbance to changes in circadian activity patterns of female chamois in summer. We expected the chamois to adjust their circadian activity pattern by avoiding being active during the hottest hours during the hottest days (H1) and during the most disturbed (i.e. weekends and public holidays) versus less disturbed (i.e. weekdays) days (H2), with a compensation of daytime activity loss at night. However, as individuals were exposed to variable levels of disturbance depending on the location of their summer home range, we also expected more marked responses in chamois that were the most exposed to hikers’ disturbance (H3), as most of them show low tolerance to hikers’ presence in this population (Courbin et al., 2022).

## MATERIALS AND METHODS

### Study Site and Species

The study took place in the National Game and Hunting Reserve of Les Bauges massif (NGHRB), a protected area in the French Alps (45°40’N, 6°13’E; 900 to 2200 m above sea level; 5200 ha). The Bauges massif is made of limestone, with cold (mean annual temperature: 9.6 °C; snow cover from November to April above 1600 m, Duparc et al., 2013) and wet (mean cumulative annual rainfall: 1600 mm, Duparc et al., 2013) climatic conditions (Appendix, Fig. A1 for details about summer temperatures). Fifty-six per cent of the area is covered by forests (treeline around 1500 m above sea level in the study site), dominated by beech, *Fagus sylvatica*, and fir, *Abies alba*, 36% by grasslands and 8% by rocks (Lopez, 2001). This heterogeneous environment leads to a huge diversity of habitats and landscapes that attracts numerous tourists every year (> 37 000, most of them during summer; 1.1 million and 0.5 million overnight stays in summer and winter, respectively, in 2014, Cloud Parc des Bauges, 2014). All alpine grasslands are crossed by a network of trails where nearly all hikers are concentrated due to the NGHRB restrictions, exposing chamois to a very localized disturbance (Courbin et al., 2022). Other recreational activities (e.g. paragliding) occur in the study site during summer, but far less frequently than hiking.

Most adult female chamois annually give birth to one young in late May. During summer, they form groups with other females, kids and yearlings that share and keep the same home range from year to year (Loison, Jullien, & Menaut, 1999). Females play a major role in the demography of the population, particularly through fecundity and survival of young females (Gaillard, Festa-Bianchet, Yoccoz, Loison, & Toigo, 2000). Chamois had no natural predators in the study area during the study period, except golden eagles, *Aquila chrysaetos*, and red foxes, *Vulpes vulpes*, which may occasionally predate newborn and injured animals. The presence of very few nonresident wolves, *Canis lupus*, has been noticed in some years (Loupfrance.fr, 2021). Hunting performed by small groups of three to four hunters occurs in the NGHRB from the beginning of September to the end of February (Courbin et al., 2022), with chamois being the main target species (ca. 100 chamois harvested per year on average, i.e. <12% of the chamois population in the area and ca. 70% of the total number of ungulates harvested), in addition to Mediterranean mouflon (ca. 20%), roe deer, *Capreolus capreolus* (ca. 4%), wild boar, *Sus scrofa* (ca. 4%) and red deer, *Cervus elaphus* (ca. 2%; see Courbin et al., 2022 for details).

### Data Collection

We focused on summers from 2004 to 2020, between 15 June and 7 September. This period includes days during and outside the French summer break which starts in early July, but excludes the hunting season which ranges from the first weekend of September to the end of February, so that chamois were exclusively exposed to hikers’ disturbance during our study period. The study period is critical for female chamois because it follows the birth season and is characterized by high energy requirements for lactation and storage of fat reserves for winter survival (Clutton-Brock, Albon, & Guinness, 1989; Richard, Toïgo, Appolinaire, Loison & Garel 2017), so that any disturbance during this period could be particularly detrimental.

We trapped 62 adult female chamois >2 years old during summer in four alpine grasslands near two trails used a lot by hikers (Armenaz and Dent des Portes). As deployment/recovery of GPS collars occurred during summer, chamois locations were not necessarily recorded daily throughout the study period. Only chamois (*N* = 62) for which we could reliably estimate the summer home range (95% kernel contour using the adehabitatHR package; Worton, 1989; Calenge, 2006) were included in the study, i.e. chamois with GPS locations available during at least two-thirds of the summer period, corresponding to 56 days in total. Animal motion was recorded every 5 min by two perpendicular captive-balls in the GPS collars monitoring neck movements (0: no movement; 255: highly mobile) and the proportion of time the animal was head down. We categorized each chamois as ‘active’ (i.e. feeding, standing, travelling or doing other activities such as interactions or scratching) or ‘inactive’ (i.e. sleeping, ruminating or resting) for every 5 min period using the discriminant model developed by Bourgoin et al. (2011; see also Darmon et al., 2014). From these animals, we obtained a total of 4464 complete days of activity records in summer (*N* = 131 chamois-years, 1285632 bouts of 5 min of activity measurements in total). The activity of each chamois was monitored during 18 - 129 days in total, corresponding to 11–49 days (mean ± SD = 34 ± 10 days, median [Q1; Q3] = 36 [26; 42] days) during a single summer period for a given chamois (Appendix, Fig. A2 and A3).

### Ethical Note

We trapped chamois using falling nets baited with salt licks. Although captures occurred during the birth and weaning period, females generally avoided drop nets (positioned on flat areas) when juveniles were particularly vulnerable (leading to females using very steep slopes as an antipredator strategy). During the weaning period, chamois form nursery groups and kids are not always closely associated with their mother, so temporary separation for periods longer than those resulting from capture/handling occur naturally in this species (Ruckstuhl & Ingold, 1998). The drop nets were triggered manually by an observer located a few hundred metres away from the nets, preventing nontarget captures, and chamois were handled immediately after the capture by several experienced people coordinated by the Office Français de la Biodiversité. As soon as an animal was handled, it was restrained with the eyes covered to reduce stress and legs tied together to avoid injuries. An average of three individuals were captured at the same time and this number was adjusted according to the number of handlers to limit the waiting time for each chamois. All efforts were made to minimize the handling time (approximately 10–15 min per individual) to release animals on site as soon as possible. Juveniles and mothers were released at the same time to prevent separation. As part of long-term capture-marking-recapture monitoring, recaptures were not avoided, although unmarked individuals were targeted when present. Since 2003, 916 individuals have been captured or recaptured of which five died (i.e. 0.55% of the individuals handled). When necessary, animals were killed by shooting a bullet in the head or by destruction of the nervous system by percussion of the cranial cavity (matador), following the ethical and legal ways recommended for large mammals after major injuries (Department of Parks and Wildlife, 2013; Legifrance, 2023). All captures, handling and sampling were conducted according to the appropriate national laws for animal welfare, following the ethical conditions detailed in the specific accreditations delivered by the Préfecture de Paris (prefectorial decree n°2009–014) in agreement with the French environmental code (Art. R421-15 to 421–31 and R422-92 to 422–94-1).

Females were equipped with GPS collars Lotek 3300S (Lotek Engineering Inc., Carp, Ontario, Canada). The GPS collars weighed < 450 g and were deployed only on individuals > 20 kg judged to be fully healthy so that it never exceeded 2.5% of the body mass of the equipped animal (average weight of females fitted with GPS collar = 30 kg, so 1.5% of individuals’ body mass on average). To allow for possible seasonal variation in neck circumference, we left a gap of ca. 3 cm between the GPS collar and the animal’s neck while making sure that the collar would not rotate. We retrieved the collar using a remote drop-off system triggered manually or automatically (TRRD or TRD drop-offs, respectively; Lotek Engineering Inc., Carp, Ontario, Canada) at the end of the monitoring of the individuals, nearly 1 year after deployment. From repeated observations of GPS-collared individuals throughout the year as part of long-term capture-mark-recapture monitoring of this population, we have no indication that GPS collars cause changes in behaviour compared to other nonequipped individuals. Similarly, we did not detect any adverse effects linked to the wearing of the collars in terms of injury or reproductive activity.

### Estimation of Variables of Interest

We assessed daily characteristics (see below) from 0139 hours UTC of day *d* to 0139 hours UTC of day *d+1*. This allowed us to explore potential compensation processes during the night following the day of interest. We selected the starting hour in accordance with the median value of the beginning of the crepuscular period in the morning during our study period. We used a microclimate model that integrates ERA5 hourly weather data produced by the Copernicus Climate Change service at ECMWF (spatial resolution 30 km x 30 km) and a digital elevation model (30 m) for downscaling atmospheric data at the population scale (micro_era5 function in NicheMapR package; Kearney & Porter, 2017; Maclean, Mosedale, & Bennie, 2019; Kearney, Gillingham, Bramer, Duffy, & Maclean, 2020). This allowed us to overcome the issue of incomplete weather data collected in the valley by the local weather station and to get standardized temperatures within the population range over the study period. Mean daily temperatures were predicted from this model at 1 m high on flat terrain for a spatial location and elevation (1950 m above sea level) corresponding to the centroid of all chamois GPS locations. We explored the effect of temperatures on female chamois activity through a categorical variable with two levels defined according to the quantiles of the mean daily temperature distribution. Days with mean temperature ≤ 25% quantile (i.e. 10.8 °C) were considered as cold days and days with mean temperature > 75% quantile (i.e. 15.5 °C) were considered as hot days. Cold days experienced by chamois ranged between 3.4 °C and 10.8 °C (mean ± SD = 8.6 ± 1.6 °C, median [Q1; Q3] = 8.9 [7.6; 10.0] °C), while hot days ranged between 15.5 °C and 20.9 °C (mean ± SD = 17.2 ± 1.2 °C, median [Q1; Q3] = 17.1 [16.2; 18.1] °C; see also Appendix, Fig. A1).

Temporal disturbance associated with hiking was defined as a categorical variable with two levels. Weekends and public holidays, i.e. vacation days such as National Day, were considered as the most disturbed days and days between Monday and Friday, other than public holidays, were considered as the least disturbed days. To check for the reliability of these two categories in the field, we compared the daily attendance of hikers (number of hikers recorded on the trail) between 0000 hours UTC and 2359 hours UTC and the hour of first arrival for both temporal disturbance levels using an eco-counter (PYRO sensor Eco-Compteur, www.eco-compteur.com; 90 cm above ground) located on Armenaz trail (45°37’18.60’’N, 6°13’2.53’’E) and present during two consecutive summers (2016 and 2017). Our study period encompassed 189 weekends and public holidays and 440 weekdays. During the same period of the year in 2016 and 2017, there were almost twice as many hikers on the trails during weekends and public holidays (mean ± SD = 74 ± 53 hikers, median [Q1; Q3] = 73 [39; 106] hikers) than during weekdays (mean = 47 ± 34 hikers, median = 40 [25; 73] hikers; Mann – Whitney *U* test: *U* = 2558, *N*_1_ = 41, *N*_2_ = 96, *P* = 0.006). Hikers also tended to arrive 60 min earlier during weekends and public holidays (mean ± SD = 0744 ± 120 min UTC, median [Q1; Q3] = 0700 [0700; 0900] hours UTC) compared to weekdays (mean ± SD = 0813 ± 180 min UTC, median [Q1; Q3] = 0800 [0700; 0900] hours UTC), although this difference was not statistically significant (Mann – Whitney *U* test: *U* = 1 559, *N*_1_ = 40, *N*_2_ = 91, *P* = 0.188; see Appendix, Fig. A4 and A5). We also conducted a Mann – Whitney *U* test to assess whether hikers’ attendance was affected by temperatures (see Results). We also ran supplementary analyses considering three contrasting temporal disturbance categories, namely (1) weekdays during the school period, (2) weekdays during the summer break and (3) weekends and public holidays (see Appendix, Section A1). These supplementary analyses provided very similar results to the original analysis contrasting two levels of disturbance (i.e. weekdays versus weekends/public holidays).

Potential spatial exposure to hikers’ disturbance was evaluated within the home range of each female chamois using attendance rates of the trails extracted from Strava Global Heatmap (Strava, 2022; www.strava.com, download date: 9 March 2022, data from the two previous years; see also Courbin et al., 2022 for a similar approach on this study site). Strava heatmap provides a value ranging from 0 for no attendance to 255 for the highest attendance for each pixel (pixel size = 25 m x 25 m) and is a good proxy of relative human attendance (see Appendix, Section A2; Thorsen et al., 2022; Venter, Gundersen, Scott & Barton, 2023). The lowest and highest values recorded for a trail pixel in our study site were 51 and 255, respectively. We thus defined attendance classes according to the quantiles of the [51; 255] distribution in order to cover trail pixels exclusively: 1 = [51; 119] (between 0% and 33% quantile, low attendance); 2 =] 119; 187] (between 33% and 66% quantiles, intermediate attendance); 3 =] 187; 255] (between 66% and 100% quantiles, high attendance). We counted trail pixels of each class in a given home range and multiplied this number by the corresponding class value (i.e. 1, 2 or 3) to give a different weight to each trail pixel according to its attendance level. We then summed these values for a given home range and normalized the total by dividing it by the home range surface to obtain our spatial disturbance index. We used the median spatial disturbance index (i.e. 0.00577) as a threshold to discriminate between most exposed chamois and least exposed chamois (see Appendix, Fig. A13).

As our study period ran over almost 3 months, daylength, which is known to influence chamois activity (Brivio et al., 2016; Fattorini et al., 2019), varied widely. We accounted for this effect by including daylength in our models (for a similar approach, see Semenzato et al., 2021). We evaluated daylength as the time elapsed between dawn and dusk using the crepuscule function from the maptools package (Bivand & Lewin-Koh, 2022; reference angle of the sun below the horizon = 6°, i.e. civil dawn/dusk).

### Statistical Analyses

We first modelled the daily activity of chamois as a series of values of either 0 (inactive) or 1 (active) for each 5 min bout using a generalized additive mixed model (GAMM) with a binomial distribution of errors (mgcv package, Wood, 2011; Wood, Goude, & Shaw 2015; Wood, 2017). We included three fixed effects in the model: (1) time of day, i.e. exact time every 5 min and (2) daylength, as well as their interaction (using the ti function dedicated to interactions in generalized additive models), as continuous variables with a cyclic cubic regression spline as smoothing basis for terms including daylength exclusively; (3) a synthesis eight-level categorical variable representing all possible combinations between temperatures classes, levels of temporal disturbance and of spatial disturbance, in interaction with the time of day (using the by argument of the s function in the implementation of the GAMM). The levels were ordered, with the reference level corresponding to a chamois living in the least exposed areas during cold weather and on weekdays. The interaction between daylength and time of day allowed us to account for the fact that the activity pattern is determined by the timing of sunrise and sunset (Ashby, 1972; Cederlund, 1989), which varies widely according to the date. We defined the unique annual id-year ID of each chamois as a random effect (using the parameter ‘re’ for the bs argument in the s function in the implementation of the GAMM) to account for interindividual variations in circadian activity pattern (see Appendix, Section A3). We set the maximum number of knots for the smoothing functions to 24. We also fitted the null model, which was similar to the model described above but excluded the temperatures-disturbance variable, for comparison purposes. All details about model fitting are provided in the Appendix, Section A4.

We used the predictions of the model to investigate the effect of temperatures and hikers’ disturbance on several activity metrics derived from this model (described below). We controlled for the effect of daylength by fixing it to the median of the complete data set (i.e. 963 min) when predicting chamois activity for each level of the temperatures-disturbance combination. We derived the total, diurnal, nocturnal and both morning and evening crepuscular activity as the area under the activity curve (AUC function of the DescTools package; Signorell, 2023) for each level of the temperatures-disturbance variable. We defined daytime as the period elapsed between 120 min after dawn and 120 min before dusk, and night-time as the period elapsed between 120 min after dusk and 120 min before dawn (see Bonnot et al., 2020 for a similar approach). Crepuscular periods corresponded to the remaining periods between daytime and night-time. We assessed the timing of the morning and evening activity peaks as the time when the highest activity probability occurs in the morning and the evening, respectively. We finally evaluated morning and evening peak duration as the period elapsed between the first time when the activity probability reached 0.5 and the last time before it went below 0.5 for both activity peaks (for a similar approach see Bourgoin et al., 2011). The percentage of variation between two situations was calculated as 100×(value_2_ – value_1_)/value_1_. We performed all analyses using the R software (R Core Development Team 2022, version R 4.3.0).

## RESULTS

### Model Adjustment

The full generalized additive mixed model (AIC = 1 578 865, explained deviance = 11.4%, adjusted *R*^2^ = 0.149) was better than the null model including time of day and daylength effects and excluding the temperatures-disturbance variable (AIC = 1 591 999, explained deviance = 10.7%, adjusted *R*^2^ = 0.140; see Appendix, Section A4) to predict activity pattern (Fig. 1). All smooth terms (fixed and random effects) were significant (*P* < 0.001).

**Figure 1:**
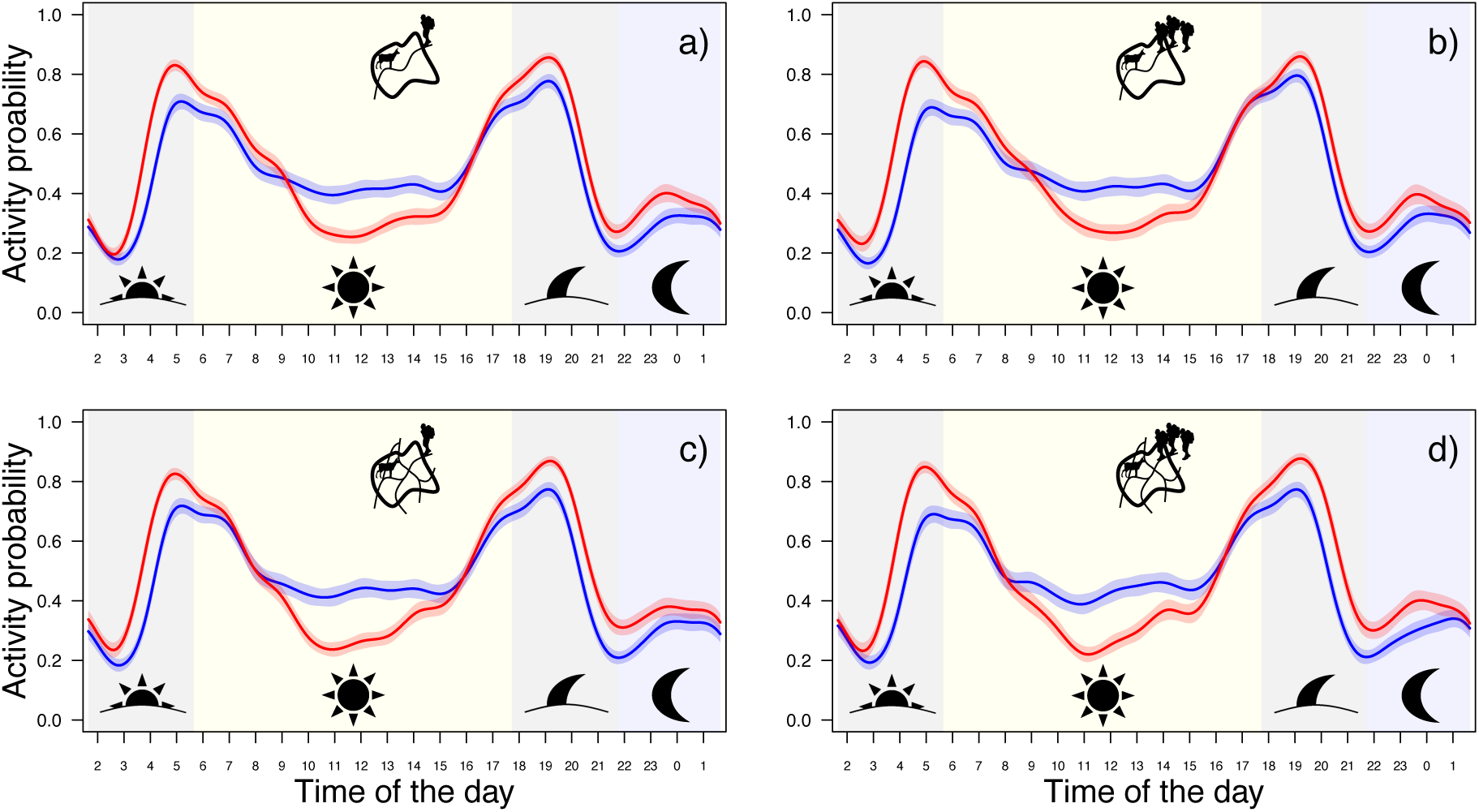
Predicted circadian activity patterns (activity ranges between 0 for inactivity to 1 for full-time activity) of 62 female northern chamois in the National Game and Hunting Reserve of Les Bauges massif, in summer (15 June to 7 September 2004–2020), according to temperatures (blue: cold day; red: hot day) and human disturbance: (a) chamois living in the least exposed areas during weekdays; (b) chamois living in the least exposed areas during weekends and public holidays; (c) chamois living in the most exposed areas during weekdays; (d) chamois living in the most exposed areas during weekends and public holidays (see also pictograms in the figure). Solid lines represent predicted values from the model and shaded areas around the curves represent 95% confidence intervals. Background shades represent the periods used to calculate the area under the activity curve for morning and evening crepuscules (grey), daytime (yellow) and night-time (blue; see also pictograms in the figure). Time of day represents hours in Coordinated Universal Time (UTC).

### Temperatures

Whatever the level of spatial exposure and temporal disturbance, chamois increased their total activity during hot days compared to cold days, by 5% on average (Fig. 2). This change corresponds to an average increase of 22% in the nocturnal activity and a decrease of 10% in the diurnal activity, from cold to hot days (Fig. 2). In addition, crepuscular activity increased from cold to hot days by more than 31% in the morning and 18% in the evening on average (Fig. 2). Chamois also advanced their morning peak by 17 min from cold to hot days, and they only slightly delayed their evening peak by 5 min on average (Fig. 3). Morning and evening peaks lasted longer during hot compared to cold days being almost 60 min and 20 min, respectively, on average (Fig. 4). Chamois maintained a certain amount of activity in the middle of the day during cold days (i.e. above 40%), whereas the activity probability at the same hours was much lower and more variable during hot days (Fig. 1).

**Figure 2:**
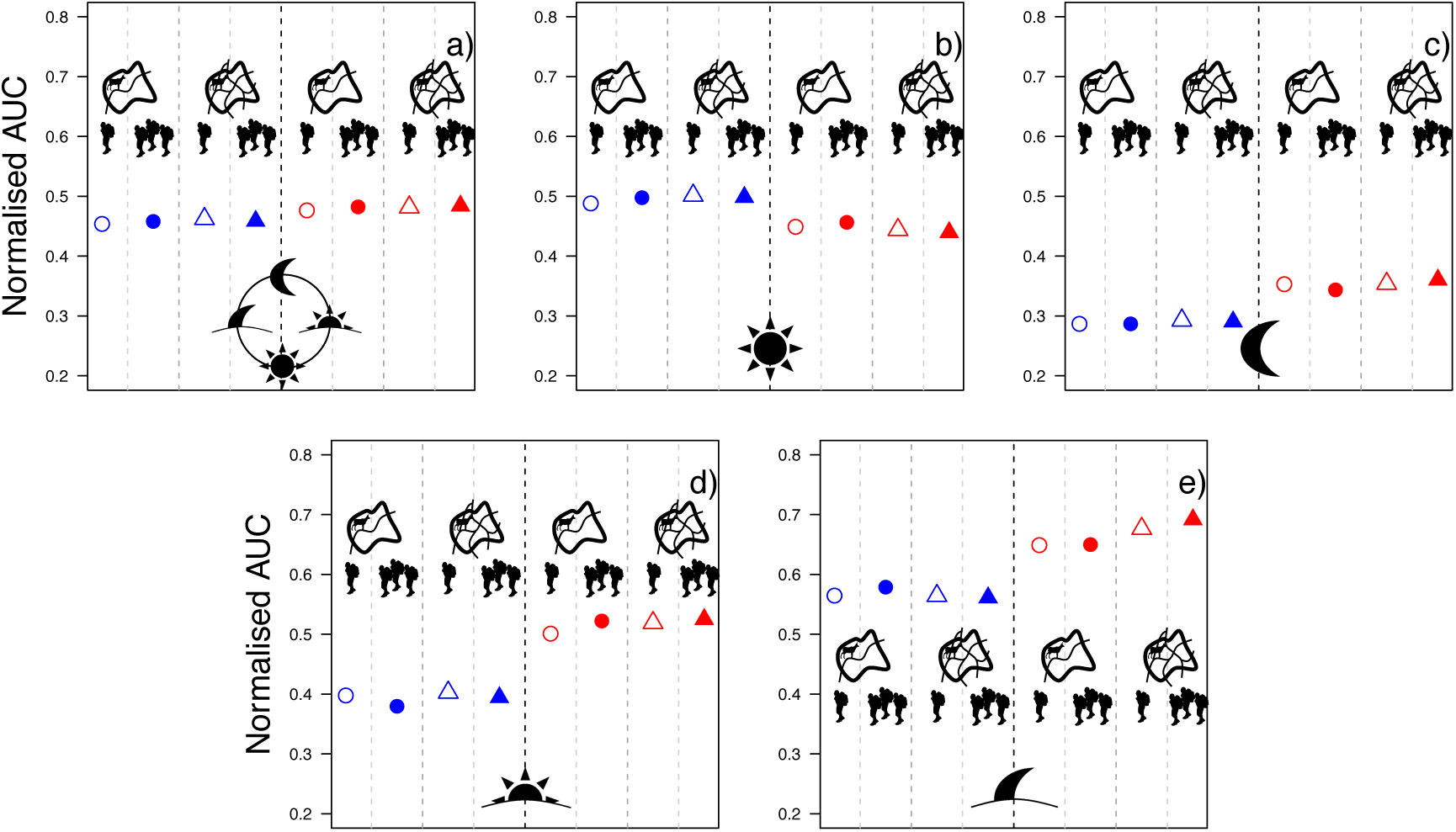
Area under curve (AUC, unitless) normalized by the duration of each period for each situation of interest, namely a combination of different levels of temperatures (blue: cold day; red: hot day), spatial disturbance(dot: low exposure; triangle: high exposure) and temporal disturbance (empty symbol: weekdays; filled symbol: weekends and public holidays; see also pictograms in the figure): (a) total (1440 min), (b) diurnal (724.8 min), (c) nocturnal (235.2 min), (d) morning crepuscular (240 min) and (e) evening crepuscular (240 min) activity (see also pictograms in the figure). AUC are calculated from predicted circadian activity patterns of northern chamois of the National Game and Hunting Reserve of Les Bauges massif, in summer (15 June to 7 September 2004–2020). We defined ‘diurnal’ as the period elapsed between 120 min after dawn and 120 min before dusk, ‘nocturnal’ as the period elapsed between 120 min after dusk and 120 min before dawn and ‘crepuscule’ as the remaining periods in the morning and evening (see text for details). Dark and light dashed lines are for visual aid to separated between the different levels of the temperatures-disturbance variable.

**Figure 3:**
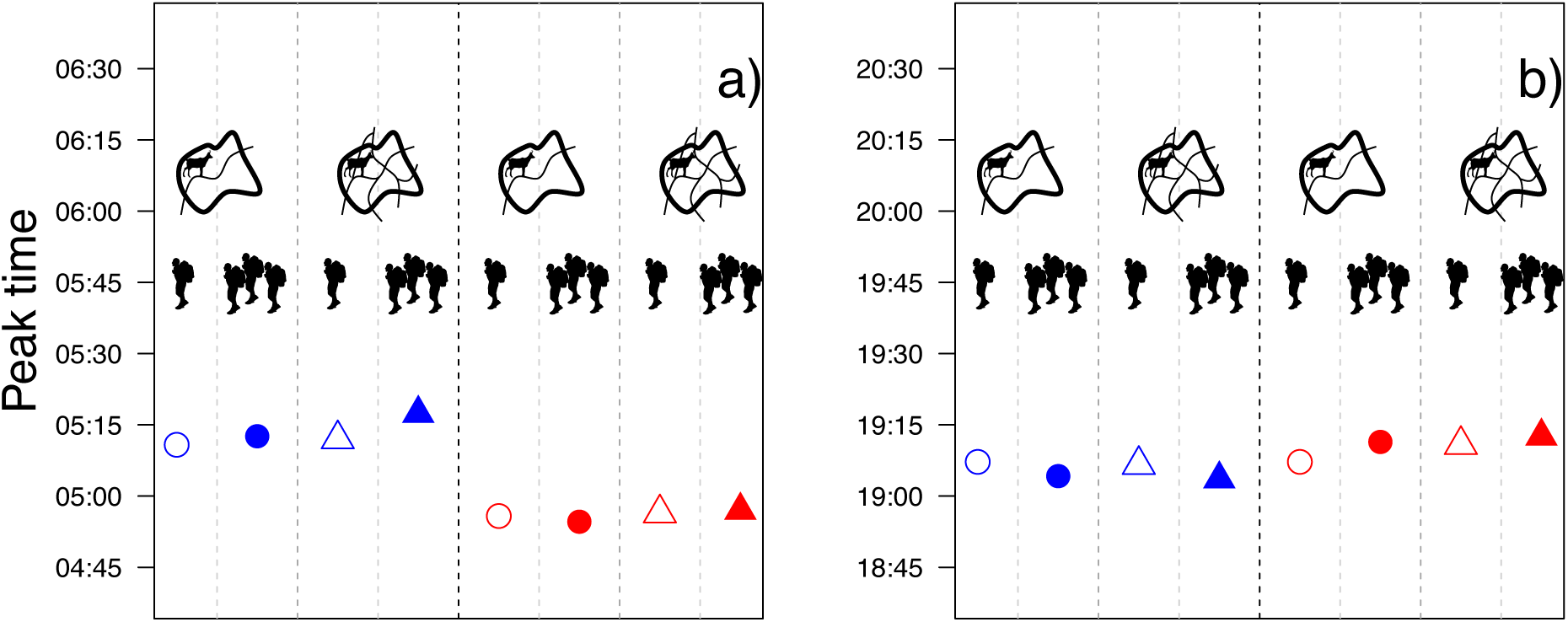
Timing of the activity peak (TAP, time of day when the activity peaks happen, i.e. the time when the highest probability of activity occurs) for each situation of interest, namely a combination of different levels of temperatures (blue: cold day; red: hot day), spatial disturbance (dot: low exposure; triangle: high exposure) and temporal disturbance (empty symbol: weekdays; filled symbol: weekends and public holidays; see also pictograms in the figure): (a) morning peak and (b) evening peak. TAP are calculated from predicted circadian activity patterns of northern chamois of National Game and Hunting Reserve of Les Bauges massif, in summer (15 June to 7 September 2004–2020). Note the change of interval on the y axis between (a) and (b), although the scale stays the same. Dark and light dashed lines are for visual aid to separated between the different levels of the temperatures-disturbance variable.

**Figure 4:**
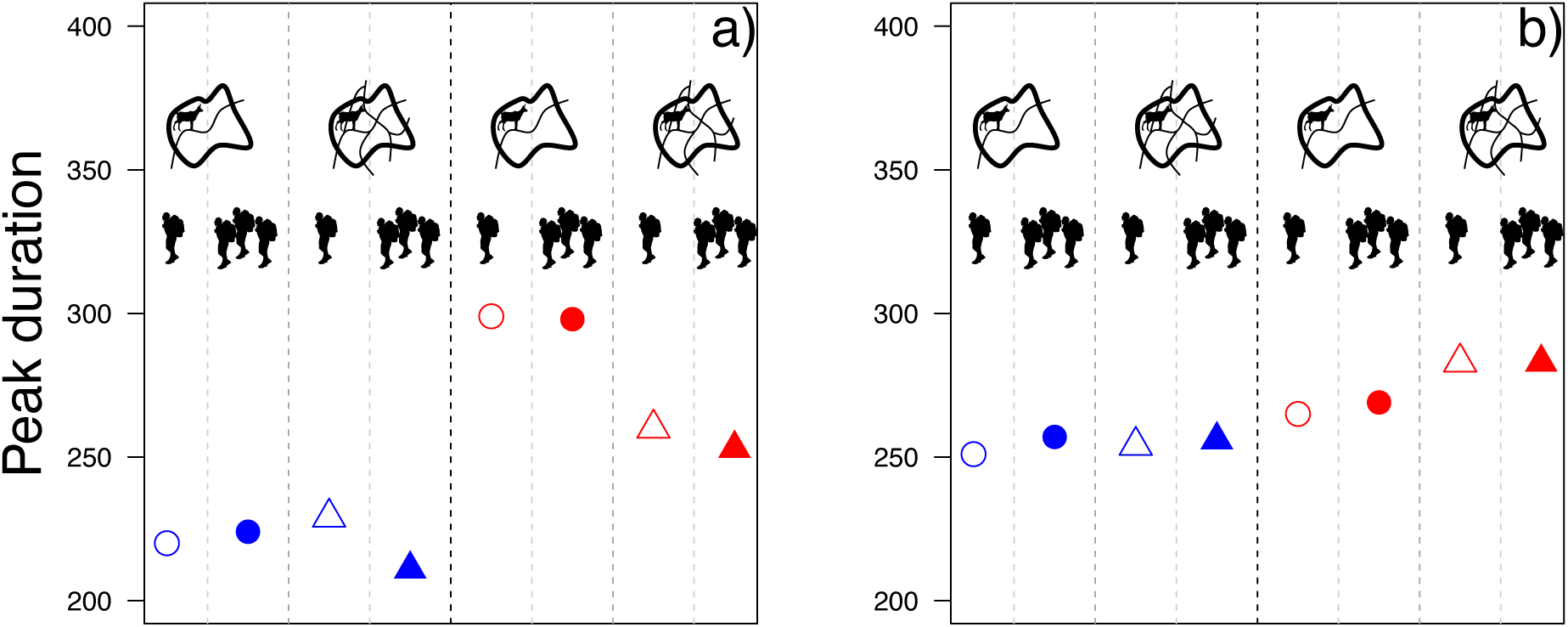
Peak duration (PD, min) for each situation of interest, namely a combination of different levels of temperatures (blue: cold day; red: hot day), spatial disturbance (dot: low exposure; triangle: high exposure) and temporal disturbance (empty symbol: weekdays; filled symbol: weekends and public holidays; see also pictograms in the figure): (a) morning and (b) evening peak. PD are calculated from predicted circadian activity patterns of northern chamois of National Game and Hunting Reserve of Les Bauges massif, in summer (15 June to 7 September 2004–2020). Dark and light dashed lines are for visual aid to separated between the different levels of the temperatures-disturbance variable.

### Hikers’ Disturbance

All the summer home ranges used by the studied chamois were crossed by at least one hiking trail. Our spatial disturbance index ranged between 0.00317 and 0.02912 (mean ± SD = 0.00699 ± 0.00433, median [Q1; Q3] = 0.00577 [0.00458; 0.00774]), with only seven chamois above 0.01000 (see Appendix, Fig. A13). In 2016 and 2017, hikers’ attendance was significantly higher when temperatures were high (mean ± SD = 45 ± 56 hikers, median [Q1; Q3] = 21 [7; 63] hikers during cold days; mean ± SD = 62 ± 37 hikers, median [Q1; Q3] = 54 [31; 84] hikers during hot days; Mann – Whitney *U* test: *U* = 333, *N*_1_ = 23, *N*_2_ = 48, *P* = 0.007).

Overall, the effect of temperature was much more pronounced than the effect of disturbance and very few differences between the general activity patterns were discernible between the different levels of hikers’ disturbance for a given temperature (Figs 1 and 5). Our analyses revealed very similar amounts of total, diurnal, nocturnal and crepuscular activity whatever the level of disturbance (Fig. 2). Similarly, the changes in terms of timing of the activity peaks according to hikers’ disturbance was not biologically significant (< 7 min; Fig. 3). Only the duration of the activity peaks during hot days (also the most disturbed days) was affected by the spatial exposure to hikers’ disturbance (Fig. 4) with the morning peak being 46 min shorter and the evening peak 20 min longer for chamois living in the most exposed areas compared to chamois living in the least exposed areas. In contrast, temporal disturbance (i.e. weekdays which are the least disturbed days versus weekends and public holidays which are the most disturbed days) had almost no effect on the length of activity peaks (Fig. 4).

**Figure 5:**
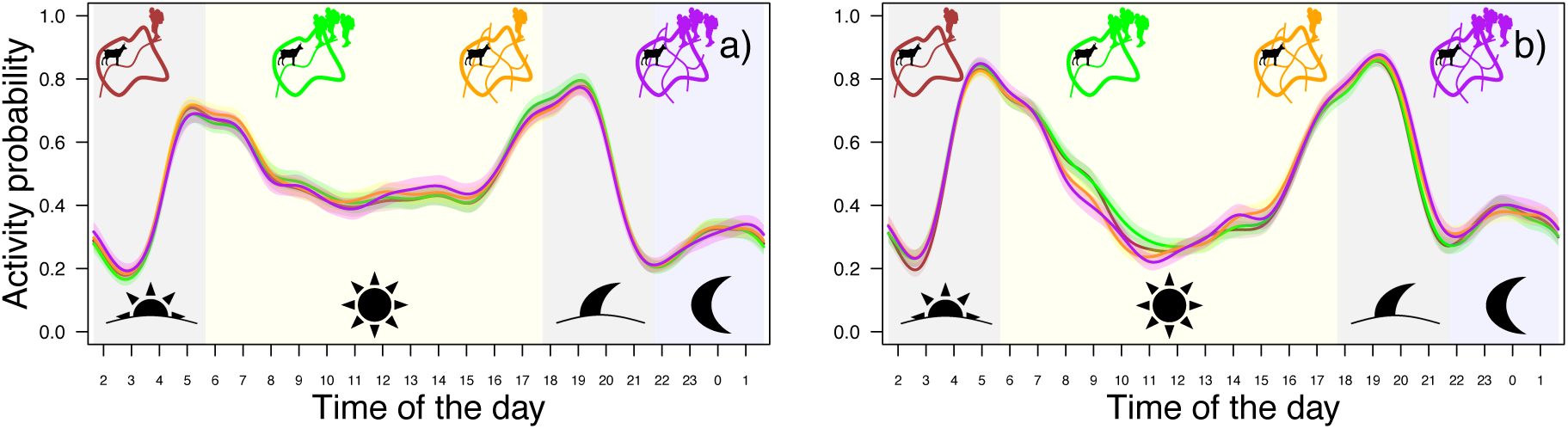
Predicted circadian activity patterns (activity ranges between 0 for inactivity and 1 for full-time activity) of female northern chamois in the National Game and Hunting Reserve of Les Bauges massif, in summer (15 June to 7 September 2004–2020), according to temperatures and human disturbance: (a) cold day and (b) hot day. In both (a) and (b), brown represents chamois living in the least exposed areas during weekdays; green represents chamois living in the least exposed areas during weekends and public holidays; orange represents chamois living in the most exposed areas during weekdays; purple represents chamois living in the most exposed areas during weekends and public holidays (see also pictograms in the figure). Solid lines represent predicted values from the model and shaded areas around the curves represent 95% confidence intervals. Background shades represent the periods used to calculate the area under the curve for morning and evening crepuscules (grey), daytime (yellow) and night-time (blue; see also pictograms in the figure). Time of day represents hours in Coordinated Universal Time (UTC).

## DISCUSSION

Our objective was to determine and compare the relative contribution of summer temperatures and hikers’ disturbance on the activity pattern of females of a mountain ungulate, the northern chamois. During hot days, chamois adjusted their circadian activity by shifting their activity peak earlier in the morning and by increasing its duration. As expected during hot days, they also increased their nocturnal and crepuscular activity while being less active during the hottest hours of the day. These behavioural adjustments allowed females to maintain a similar level of total daily activity between cold and hot days. On the other hand, and contrary to our expectations, chamois only rarely adjusted their activity pattern according to hikers’ disturbance (see also Appendix, Section A1). Only during hot days (also the most disturbed days) did we observe a marked decrease in the duration of the morning activity peak and a slight increase in the duration of the evening activity peak for chamois living in the most exposed areas compared to chamois living in the least exposed areas, as well as a slight increase in the nocturnal and crepuscular activity at the expense of the diurnal activity. Temporal disturbance (i.e. weekends and public holidays versus weekdays) had almost no effect on chamois activity peaks.

Although chamois seem to be less sensitive to high temperatures than other alpine ungulate species (see Darmon et al., 2014 for a comparison between chamois and mouflon; see also Semenzato et al. 2021 for Alpine ibex), their faecal glucocorticoid metabolite concentration increases during drought conditions in summer (Anderwald, Campell Andri, & Palme 2021), as well as the time they spend in thermal refuges during hot days (Malagnino, 2022). This may explain why chamois responded much more strongly to temperatures than to hikers’ disturbance. Moreover, in our study site, hikers’ attendance was significantly higher when temperatures were high. Hence, the behavioural changes made to buffer chamois from high temperatures may also protect them from hikers’ disturbance, as both factors were related. Animals have to face several stressors (e.g. human disturbance, predation, high temperatures) and they cannot indefinitely adjust their behavioural response as stressors accumulate (Benoist, Garel, Cugnasse, & Blanchard, 2013). For example, during cooler periods, yellow-blotched sawback, *Graptemys flavimaculata*, tend to increase their basking time even when disturbed by humans, presumably because low temperatures force them to favour thermoregulation at the expense of risk avoidance (Selman, Qualls, & Owen, 2013). Moreover, most of the response to human disturbance in our chamois population seemed to happen at the spatial level rather than the temporal level (Fig. 4a) and only during the hot days (when hikers’ attendance was at its highest level), causing a marked decline in the duration of the morning activity peak in chamois living in the most versus least exposed areas.

Recent studies have provided evidence of a non-negligeable nocturnal activity peak in chamois (Carnevali, Lovari, Monaco, & Mori, 2016). Here, we also highlighted that nocturnal activity significantly increased while diurnal activity decreased during hot days, confirming the influence of high temperatures on chamois activity. As in numerous other species, chamois decreased their diurnal activity when exposed to high temperatures and/or human disturbance and attempted to compensate for the activity loss when they were absent (e.g. later in the day or at night, Gaynor, Hojnowski, Carter, & Brashares, 2018). Similarly, the lynx, *Lynx lynx*, responds spatially to disturbance by using the areas located close to the trails inside its home range less frequently during the day than at night (Thorsen et al., 2022). However, an increase in nocturnal activity could expose individuals to other threats, such as nocturnal predators (Bonnot et al., 2020). This is particularly true in our study site with the progressive return of wolves recorded since 2020. Large herbivores may face the difficult challenge of avoiding being exposed to high temperatures while limiting their exposure to human disturbance, but without shifting completely their activity at night because of predation risk. The shift to more crepuscular activity observed in this study could be an answer to this trade-off.

Interestingly and in contrast with Brivio et al. (2016) and Grignolio et al. (2018) in another chamois population, total activity was slightly higher (+5%) during hot than cold days in our chamois population. Such a finding is not necessarily surprising as other studies have found similar results (in the same population, Malagnino, 2022; in Alpine ibex maintaining a constant activity both during cold and hot days, Semenzato et al., 2021). In addition, this does not necessarily translate into an increase in feeding behaviour but might be because chamois display diel migrations between refuges and feeding spots (Courbin et al., 2022), particularly if both habitats are distant or heterogeneous. With the increasing temperatures in summer, chamois could be forced to commute more frequently or move further to find shelter from the heat while maintaining the same level of feeding activity, eventually leading to the overall increase observed. Unfortunately, our data provide no clue about the nature of the behaviours displayed (i.e. foraging, moving or social behaviours) and we cannot ascertain whether this rise in activity effectively corresponds to spatial displacement. Likewise, the pigmy rabbit, *Brachylagus idahoensis*, selects significantly more ground depressions as refuges during summer to mitigate the adverse effects of high temperatures (Milling, Rachlow, Johnson, Forbey, & Shipley, 2017).

We observed only minor differences between the most and least disturbed chamois, either because chamois respond similarly to human presence regardless of the encounter risk because hikers are present almost every day even if at different levels (especially during summer with the summer school break), or because they shift their activity in habitats further away from the main source of disturbance, i.e. the trails (Courbin et al. 2022). In our population, 85% of the individuals temporarily move away from trails every day when hikers are present (Courbin et al., 2022), demonstrating that even the less exposed chamois adjust their behaviour to human disturbance. This low tolerance affects the whole population and not just the most exposed individuals and might explain the weak difference observed between groups. Wolves also display high sensitivity to human disturbance and completely avoid the areas frequented by tourists, even if the level of attendance is as low as 40 visitors per week (Sytsma, Lewis, Gardner, & Prugh, 2022).

Mountain animals face both rising temperatures from ongoing climate warming, and modifications in their exposure to actual and perceived risks from the development of recreational activities and the return of predators in these landscapes. The interaction between those factors could eventually jeopardise their demography if particular attention is not paid to conserving habitats providing both refuge against summer heat and human disturbance (Frid & Dill, 2002; Chirichella, Stephens, Mason, & Apollonio, 2021). Moreover, both constraints can lead either to similar responses (e.g. rest in refuge areas), or to opposite reactions (e.g. reduce activity to avoid thermal stress, increase movement to reach thermal refuges or escape to limit predation risk), which complicates our understanding of their behavioural response. Here we highlight a strong effect of temperatures but low effects of hikers’ disturbance on the activity pattern of northern female chamois. Identifying the effect of human disturbance on wildlife remains a difficult task, particularly at the population scale (Tablado & Jenni, 2017), and an integrated approach is needed to consider all forms of response, such as variations in activity level and timing, behavioural budgets and spatial displacements.

## Author Contributions

**Lucie Thel**: Conceptualization; Data curation; Formal analysis; Investigation; Methodology; Visualization; Writing–Original draft; Writing–Review & editing. **Mathieu Garel**: Conceptualization; Data curation; Formal analysis; Funding acquisition; Investigation; Methodology; Supervision; Validation; Visualization; Writing–Review & editing. **Pascal Marchand**: Conceptualization; Data curation; Methodology; Supervision; Validation; Writing–Review & editing. **Gilles Bourgoin**: Conceptualization; Methodology; Supervision; Validation; Writing–Review & editing. **Anne Loison**: Conceptualization; Funding acquisition; Methodology; Project administration; Resources; Supervision; Validation; Writing–Review & editing.

## Data Availability

Data are available from the Zenodo Repository: https://zenodo.org/record/8123239.

## Acknowledgments

This study was funded by Agence Nationale de la Recherche (French National Research Agency, ANR) Grant Mov-It No.16-CE02-0010 coordinated by Anne Loison, Centre National de la Recherche Scientifique (Centre National de la Recherche Scientifique, CNRS). We thank all participants for helping with the capture and marking of animals, for the recovery of GPS collars, and for collecting GPS data on human activities: volunteers and professionals from the Office Français de la Biodiversité (formerly Office National de la Chasse et de la Faune Sauvage), the Office National des Forêts, the Groupement d’Intérêt Cynégétique des Bauges, the Parc Naturel Régional du massif des Bauges, the Observatoire Grande Faune et Habitats and the laboratory Environnements Dynamiques Territoires Montagnes (EDYTEM). This work was performed using the computing facilities of the IFB Cloud.

## APPENDIX

### Section A1: Complementary analyses of the effect of temperature and hikers disturbance on chamois activity patterns using three levels of temporal disturbance

We ran supplementary analyses considering three contrasting temporal disturbance categories, namely (1) weekdays during the school period, (2) weekdays during the summer break and (3) weekends and public holidays. These supplementary analyses provided very similar results to the original analysis contrasting two levels of disturbance (i.e. weekdays versus weekends/public holidays, see main text). Compare Fig. 1 versus A6, Fig. 2 versus A7, Fig. 3 versus A8, Fig. 4 versus A9 and Fig. 5 versus A10.

### Section A2: Comparison Between Strava Map and GPS Data

GPS trackers were distributed to hikers’ groups during summer 2014 and 2015 in two sites of the National Game and Hunting Reserve of Les Bauges massif (NGHRB; Armenaz trail: *N*= 73 groups; Dent des Portes trail: *N* = 114 groups) to study space use by hikers (Kerouanton, 2020). Trails were segmented according to the number of people using them (*N* = 62 segments) and we evaluated the proportion of use for each trail section, relative to each study site. This protocol allowed us to precisely monitor hikers’ spatial and temporal behaviour in our study site, but (1) only a very small area of the study site was covered, (2) during a short period of time which solely encompassed a small part of our study period (2004–2020), and (3) illustrated the spatial behaviour of a small number of hikers only.

Strava (www.strava.com) is a training app to which trail runners and hikers can voluntarily upload their GPS tracks. Strava Global Heatmap is a downloadable heatmap which illustrates the attendance rates of the trails via values ranging from 0 for no attendance to 255 for the highest attendance for each pixel of the map (Strava, 2022). In the NGHRB, the lowest value recorded for a trail pixel was 51 and the highest was 255. The main advantage of Strava consists in its high precision and large coverage to capture spatial and temporal variation in recreational activities, although Strava users only represent a low proportion and a nonrandom sample of the recreational population (Venter, Gundersen, Scott & Barton, 2023).

#### Methods

First, we compared track overlap by visual inspection between Strava and GPS data sets in the NGHRB. Second, we compared the relative attendance by trail runners and hikers assessed from Strava and GPS data sets, respectively. We extracted the mean value of all the pixels of the Strava map located on a given trail segment of the GPS map. We tested for association between this mean value and the relative proportion of use of the corresponding segment with the Pearson correlation test.

#### Results and discussion

Strava and GPS trails (where available) overlapped very well (Fig. A11). Some trails used by the hikers equipped with GPS trackers in Armenaz sector seemed to be only rarely used by trail runners according to the Strava map. Nevertheless, both attendance measures were highly correlated (Pearson correlation: ⍴ = 0.55, *r* = 3.5, *N* = 61, *P* = 0.002; Fig. A12). We concluded that Strava is a good proxy of relative human attendance, in particular hikers’ attendance, in our study site, and can be used to predict hikers’ attendance at a large spatial scale even in the absence of GPS tracking of hikers (see also Courbin et al., 2022).

### Section A3: Interindividual Variability in the Circadian Activity Pattern in Chamois

All the chamois in the study displayed the characteristic bimodal activity pattern (i.e. one activity peak in the early morning and one in the evening), with a smaller peak during the night, as illustrated by the mean activity pattern calculated from all chamois and all days available (raw data, Fig. A14). However, the circadian activity patterns of the chamois were also characterized by a high interindividual variability. The amplitude (compare Fig. A15a and b) and the spread (compare Fig. A15a and c) of the activity peaks varied widely among individuals. Several chamois displayed a small peak during the day (e.g. Fig. A15d). These variations could be due to individual personality, age (Watanuki & Nakayama, 1993), different life histories such as the presence or absence of a kid (Lesmerises, Déry, Johnson, & St-Laurent, 2017; but most adult females give birth in the site) or external conditions experienced by the individual in its home range (Bourgoin et al., 2011). It could also be due to data collection. Each chamois was equipped with a GPS collar and an activity sensor, whose sensitivity to movement can vary intrinsically and according to weather conditions experienced by the chamois, among other factors. Moreover, collars were put on and removed between June and September, which is also the period of interest in our study. Thus, the number of monitored days was highly variable among chamois (Fig. A3).

### Section A4: Fitting a Generalized Additive Mixed Model to Describe Activity Patterns

We adjusted a generalized additive mixed model with a binomial distribution of errors to our activity data (northern chamois in the National Game and Hunting Reserve of Les Bauges massif, 2004–2020) with the maximum number of knots *k* = 24 to predict the activity pattern for each of the eight levels of the temperatures-disturbance categorical variable for a theoretical chamois (i.e. random effect set to null). The procedure is described in the Methods and the relevant outputs of the model are presented in the Results. Tables A1, A2, A3, A4 and A5 present the raw outputs from the summary function (model outputs, approximate significance of smooth terms) and the gam.check function (basis dimension checking; mgcv package, Wood2011, Wood2015, Wood2017) calculated with R software (R Core Development Team 2022, version R 4.3.0).

### Tables

**Table A1:** Median of the total number of hikers per day on the Armenaz trail in the National Game and Hunting Reserve of Les Bauges massif (part of summer 2016 and summer 2017)

| Day | Median of the total number of<br>hikers on the trail | Relative difference (%) as compared to<br>the least disturbed day (i.e. Tuesday) |
| --- | --- | --- |
| Monday | 44 | + 42 |
| Tuesday | 31 | 0 |
| Wednesday | 39 | + 26 |
| Thursday | 53 | + 71 |
| Friday | 39 | + 26 |
| **Saturday** | **70** | **+ 126** |
| **Sunday** | **83** | **+ 168** |
| **Holiday** | **114** | **+ 268** |

\*Days in bold type correspond to weekends and public holidays.

**Table A2:** Sample size for each level of the synthesis eight-levels categorical variable representing all possible combinations between temperatures classes, levels of temporal.

| Temperatures-disturbance variable levels |  |  | Sample size<br>(total no. of<br>activity data) | No. of<br>chamois-<br>days |
| --- | --- | --- | --- | --- |
| Spatial<br>disturbance | Temporal disturbance | Temperatures |  |  |
| Low | Weekdays | Low | 244 224 | 848 |
| Low | Weekends & public | Low | 120 096 | 417 |
| Low | Weekdays | High | 241 632 | 839 |
| Low | Weekends & public | High | 87 264 | 303 |
| High | Weekdays | Low | 213 984 | 743 |
| High | Weekends & public | Low | 104 832 | 364 |
| High | Weekdays | High | 202 752 | 704 |
| High | Weekends & public | High | 70 848 | 246 |

**Table A3:** Model outputs as presented by the summary function of R software (R Core Development Team 2022, version R 4.3.0)

|  | Estimate | SE | Z | Pr (>\|z\|) |
| --- | --- | --- | --- | --- |
| (Intercept) | -0.077445 | 0.028887 | -2.681 | 0.007** |
| Three.L | 0.167552 | 0.036227 | 4.625 | 3.74e-06*** |
| Three.Q | 0.003719 | 0.006633 | 0.561 | 0.575 |
| Three.C | -0.033791 | 0.043036 | -0.785 | 0.432 |
| Three^4 | -0.010713 | 0.005927 | -1.808 | 0.071 |
| Three^5 | 0.031552 | 0.059470 | 0.531 | 0.596 |
| Three^6 | 0.019075 | 0.006381 | 2.989 | 0.003** |
| Three^7 | -0.029235 | 0.010000 | -2.923 | 0.003** |

Family: binomial; link function: logit; formula: actcod~three+s(heure\_num2, bs='cc', k=24)+s(heure\_num2, bs='cc', k=24, by=three, m=1)+s(length\_day\_min, k=24)+ti(heure\_num2, length\_day\_min, bs='cc' , k=24)+s(id\_year, bs='re'); signif. $R^2$ adj=0.149; deviance explained=11.4%; fREML=1.8258e+06; scale estimate=1; N=1 285 632. \*\*\* $P < 0.001$ ; \*\* $P < 0.01$ .

In the table, “actcod” corresponds to the activity of chamois (0: inactive or 1: active), “three” corresponds to the temperatures-disturbance variable, “heure_num2” corresponds to time of day variable, “length_day_min” corresponds to day length variable, “id_year” corresponds to the year.

**Table A4:**
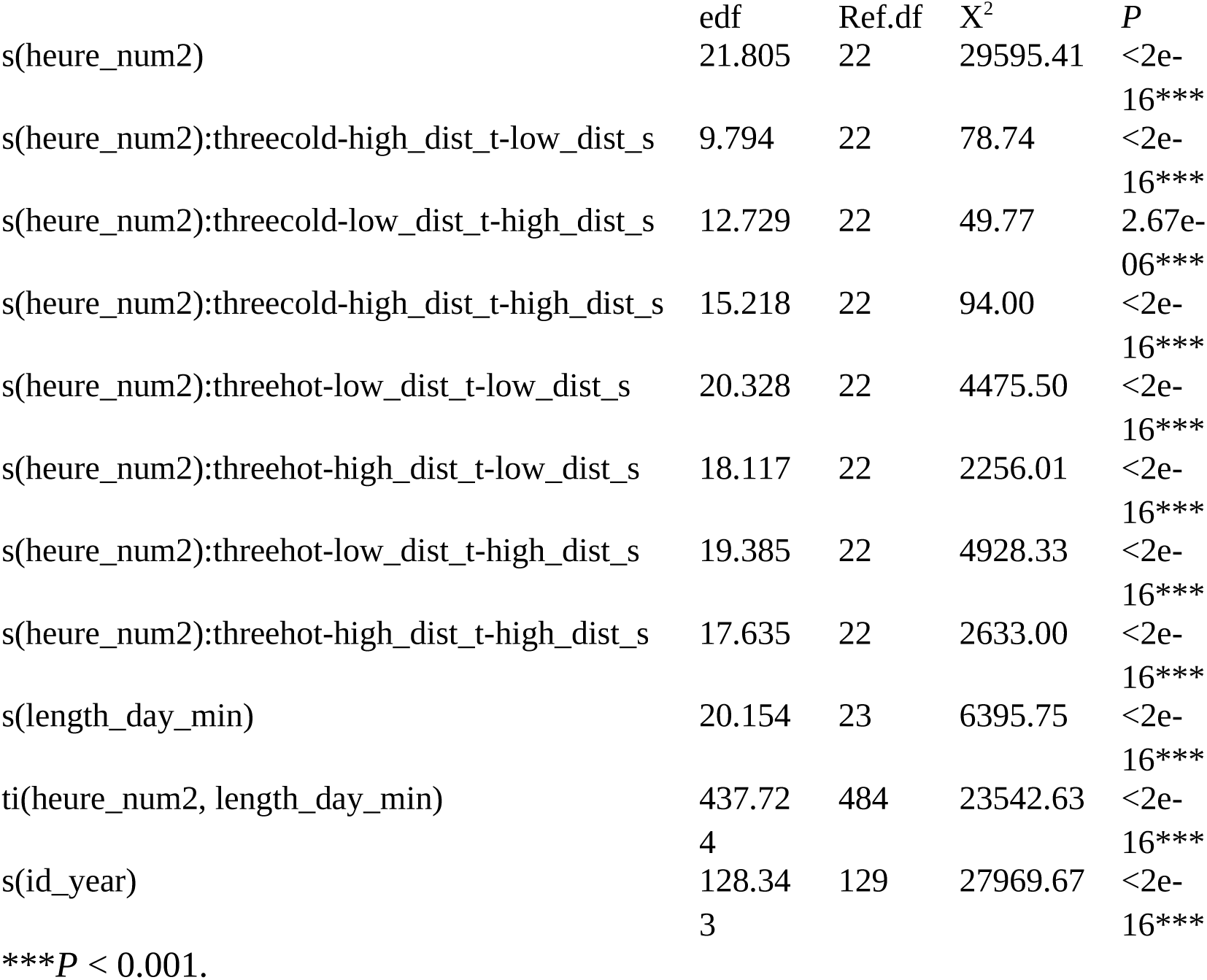
checking of approximate significance of smooth terms (extract from the outputs of the summary function)

In the table, “heure_num2” corresponds to time of day variable, “length_day_min” corresponds to day length variable, “id_year” corresponds to the year. “cold” and “hot” correspond to the two levels of the temperature variable. “dist_t” and “dist_s” correspond to the temporal and spatial disturbance variables respectively. These variables can be either “low” or “high”. For instance, threecold-high_dist_t-low_dist_s corresponds to the level of the temperatures-disturbance variable of a cold day, which is also a weekend or public holiday, for a chamois living in the least exposed areas.

**Table A5:**
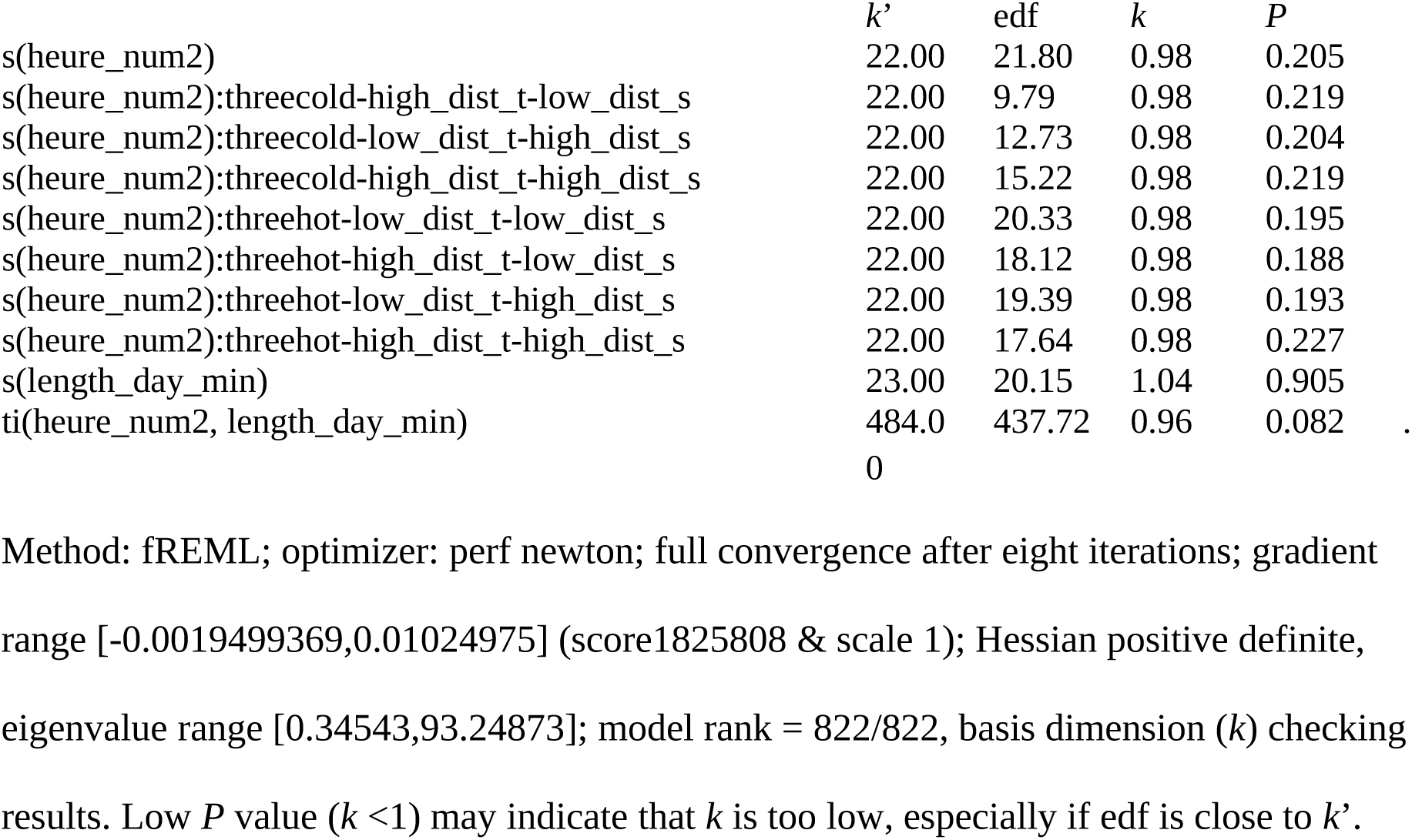
Checking of the basis dimension (knots) using the gam.check function.

In the table, “heure_num2” corresponds to time of day variable, “length_day_min” corresponds to day length variable, “id_year” corresponds to the year. “cold” and “hot” correspond to the two levels of the temperature variable. “dist_t” and “dist_s” correspond to the temporal and spatial disturbance variables respectively. These variables can be either “low” or “high”. For instance, threecold-high_dist_t-low_dist_s corresponds to the level of the temperatures-disturbance variable of a cold day, which is also a weekend or public holiday, for a chamois living in the least exposed areas.

**Figure A1:**
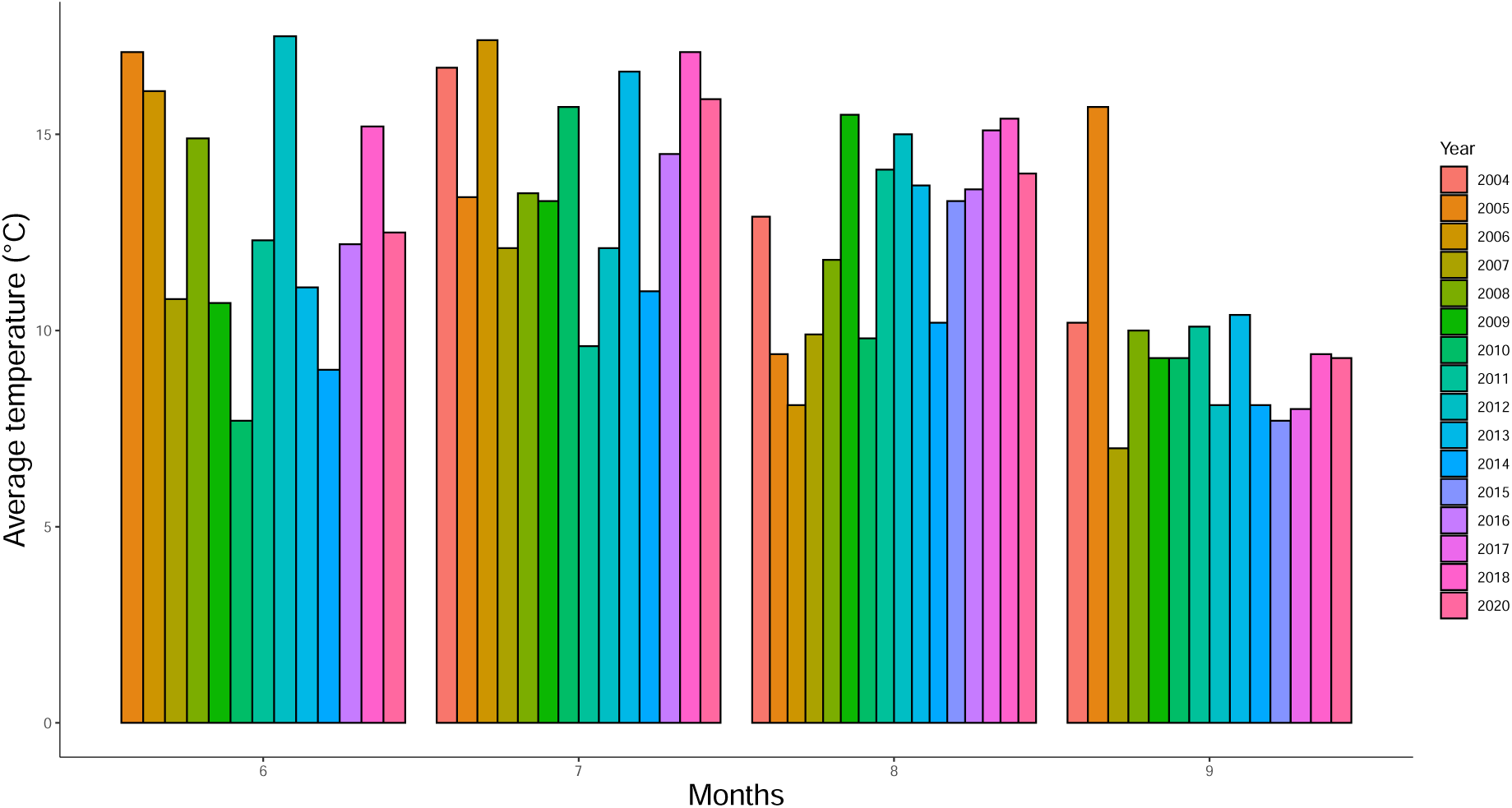
Monthly mean of daily temperatures (°C) in summer over the study period (between 2004 and 2020). Months: 6 = June, 7 = July, 8 = August, 9 = September.

**Figure A2:**
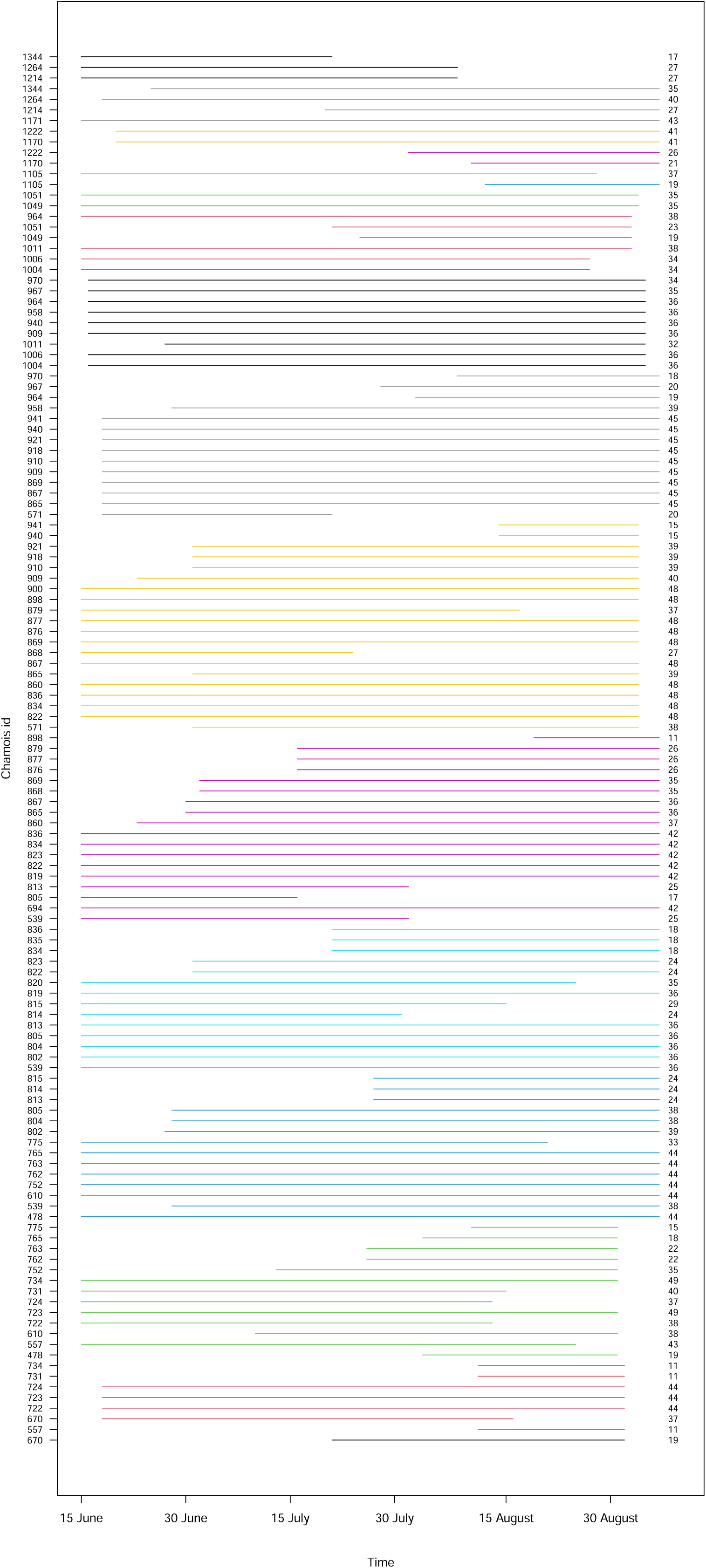
Sampling period for each collared female northern chamois (*N* = 62) every year (represented by the different colours, from bottom, i.e. 2004 to top, i.e. 2020) in the National Game and Hunting Reserve of Les Bauges massif, in summer (15 June to 7 September). The numbers on the left of each sampling period correspond to the chamois identity and can be repeated more than once as some chamois were collared during several years. The numbers on the right of each sampling period correspond to the number of days included in the analyses, i.e. when the activity monitoring was complete over the 1440 min period (i.e. 24 h; 288 records, i.e. one record for every 5 min bout).

**Figure A3:**
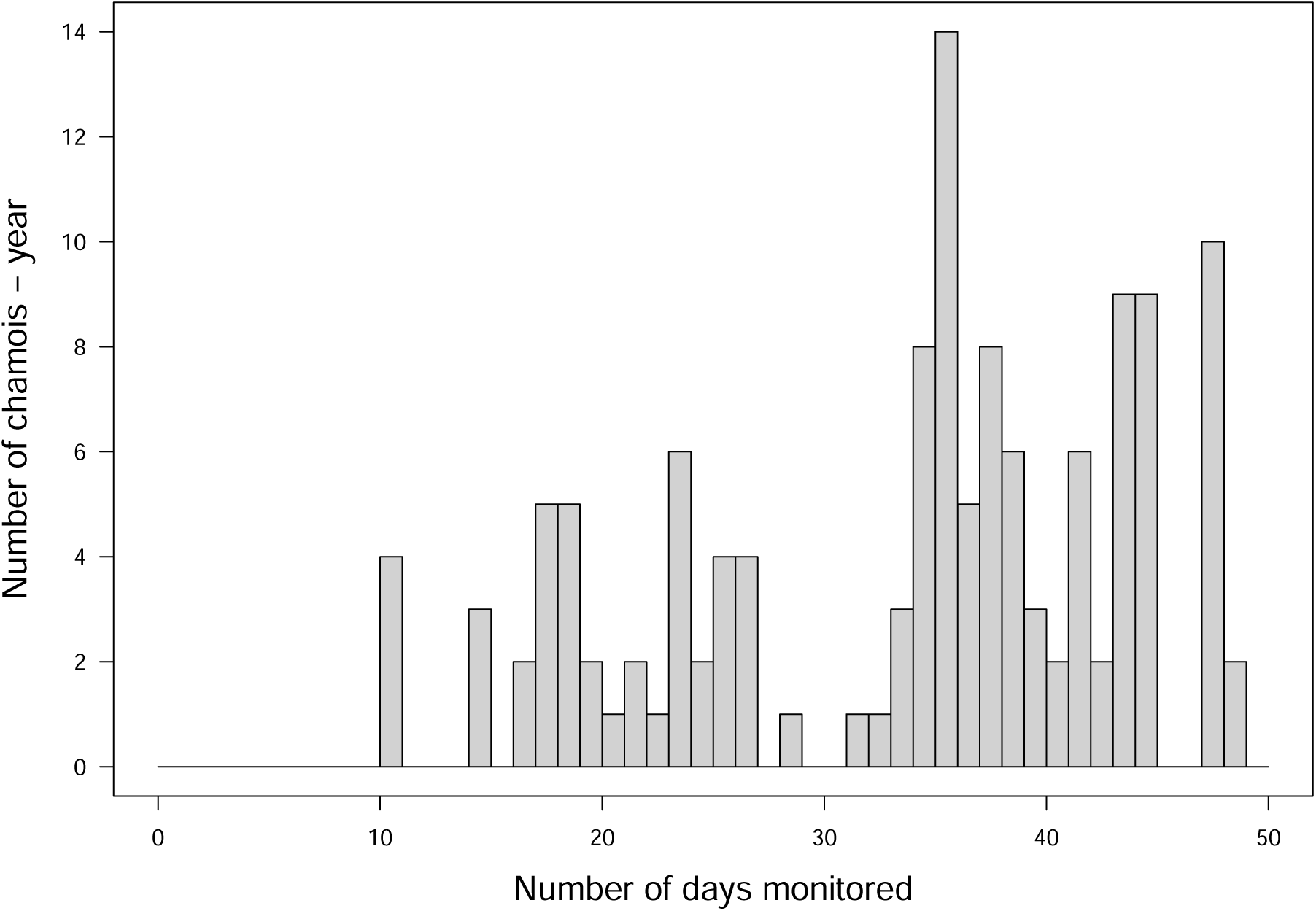
Distribution of chamois according to the number of days they were monitored during one summer (*N* = 131 chamois-years, chamois could be monitored during several consecutive summers) in the National Game and Hunting Reserve of Les Bauges massif between 2004 and 2020 (15 June to 7 September).

**Figure A4:**
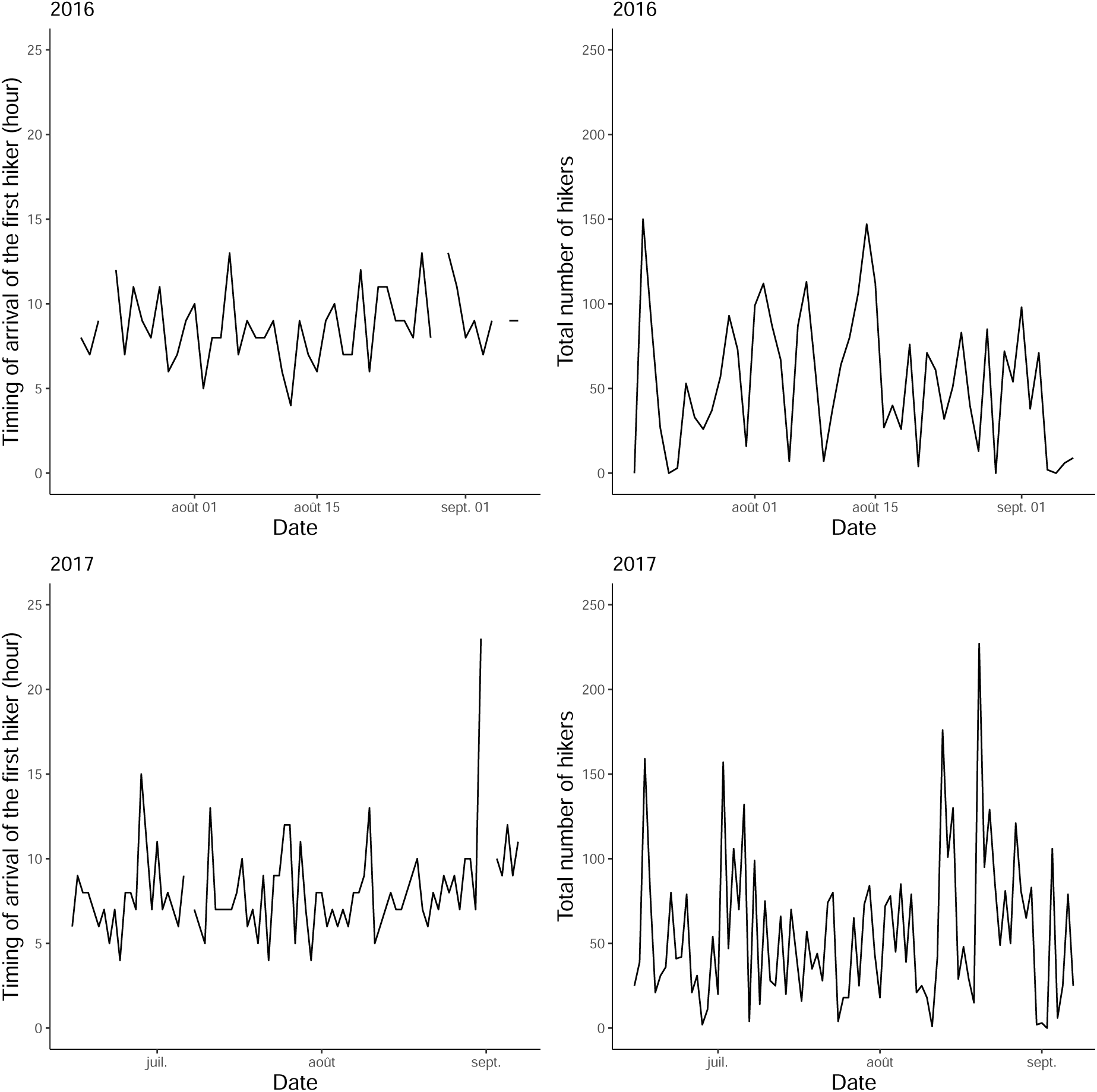
Daily hikers’ attendance as the time of arrival of the first hiker going up the mountain and the total number of hikers during the day (between 0000 hours UTC and 2359 hours UTC, part of summer 2016 and summer 2017).

**Figure A5:**
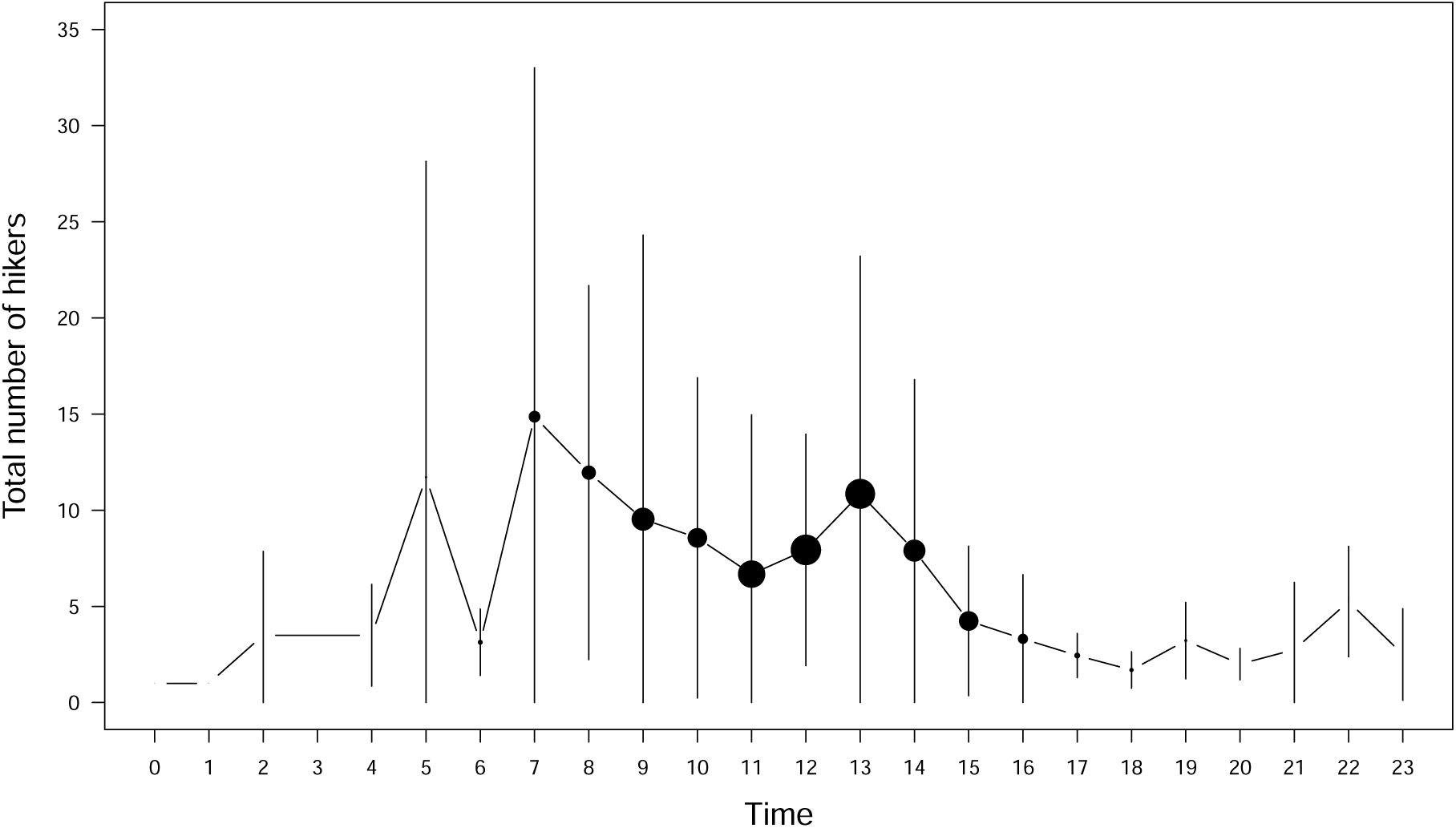
Mean total number of hikers present on the trails (part of summer 2016 and summer 2017) in the study site (National Game and Hunting Reserve of Les Bauges massif) according to the hour of the day (in hours, Coordinated Universal Time UTC). The size of each point is proportional to the number of days when at least one hiker was seen for this hour. The vertical bars correspond to the standard deviation. As the counter is located at the beginning of the trail, hikers are expected to reach the chamois area approximately 1 h after their presence has been detected.

**Figure A6:**
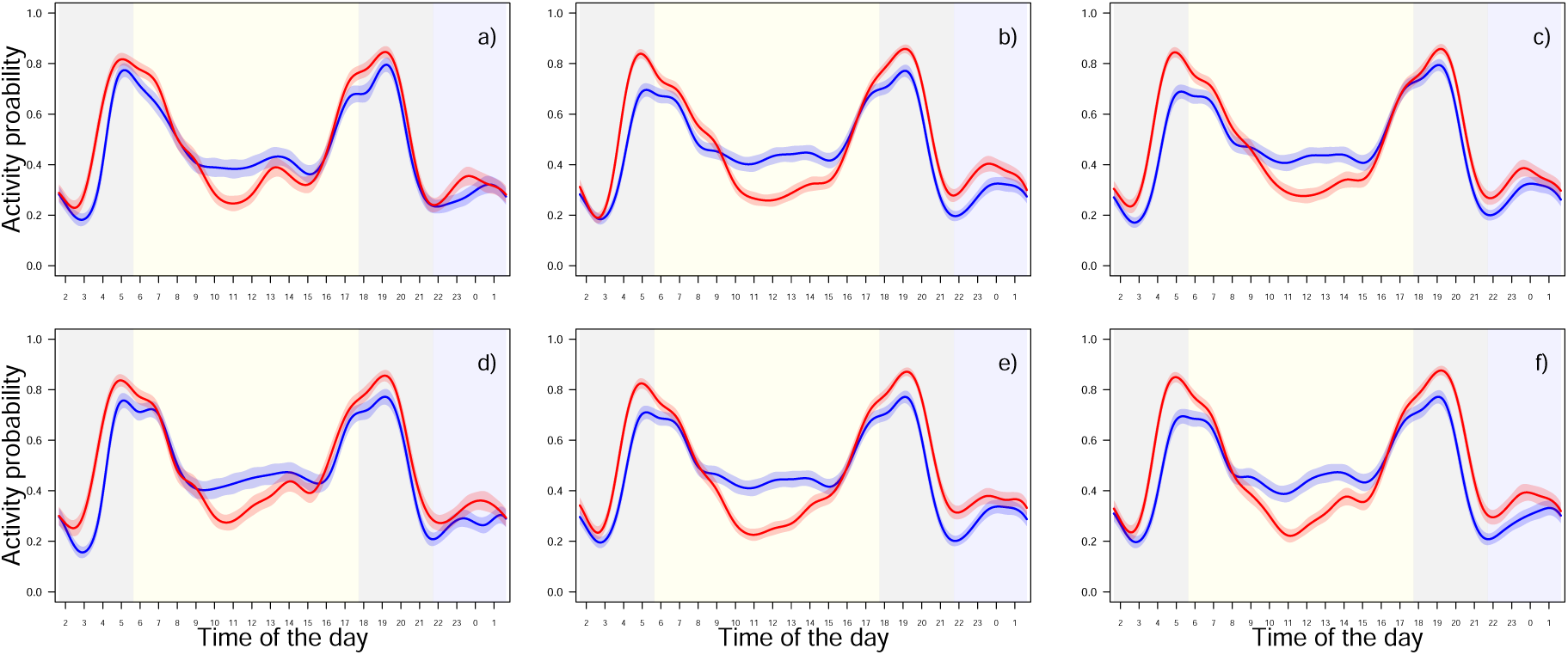
Predicted circadian activity patterns (activity ranges between 0 for inactivity and 1 for full-time activity) of 62 female northern chamois in the National Game and Hunting Reserve of Les Bauges massif, in summer (15 June to 7 September 2004–2020), according to temperatures (blue: cold day; red: hot day) and human disturbance: (a) chamois living in the least exposed areas during weekdays of the school period; (b) chamois living in the least exposed areas during weekdays of the summer break; (c) chamois living in the least exposed areas during weekends and public holidays; (d) chamois living in the most exposed areas during weekdays of the school period; (e) chamois living in the most exposed areas during weekdays of the summer break; (f) chamois living in the most exposed areas during weekends and public holidays. Solid lines represent predicted values from the model and shaded areas around the curves represent 95% confidence intervals. Background shades represent the periods used to calculate the area under the activity curve for morning and evening crepuscules (grey) and daytime (yellow). Time of day represents hours in Coordinated Universal Time (UTC).

**Figure A7:**
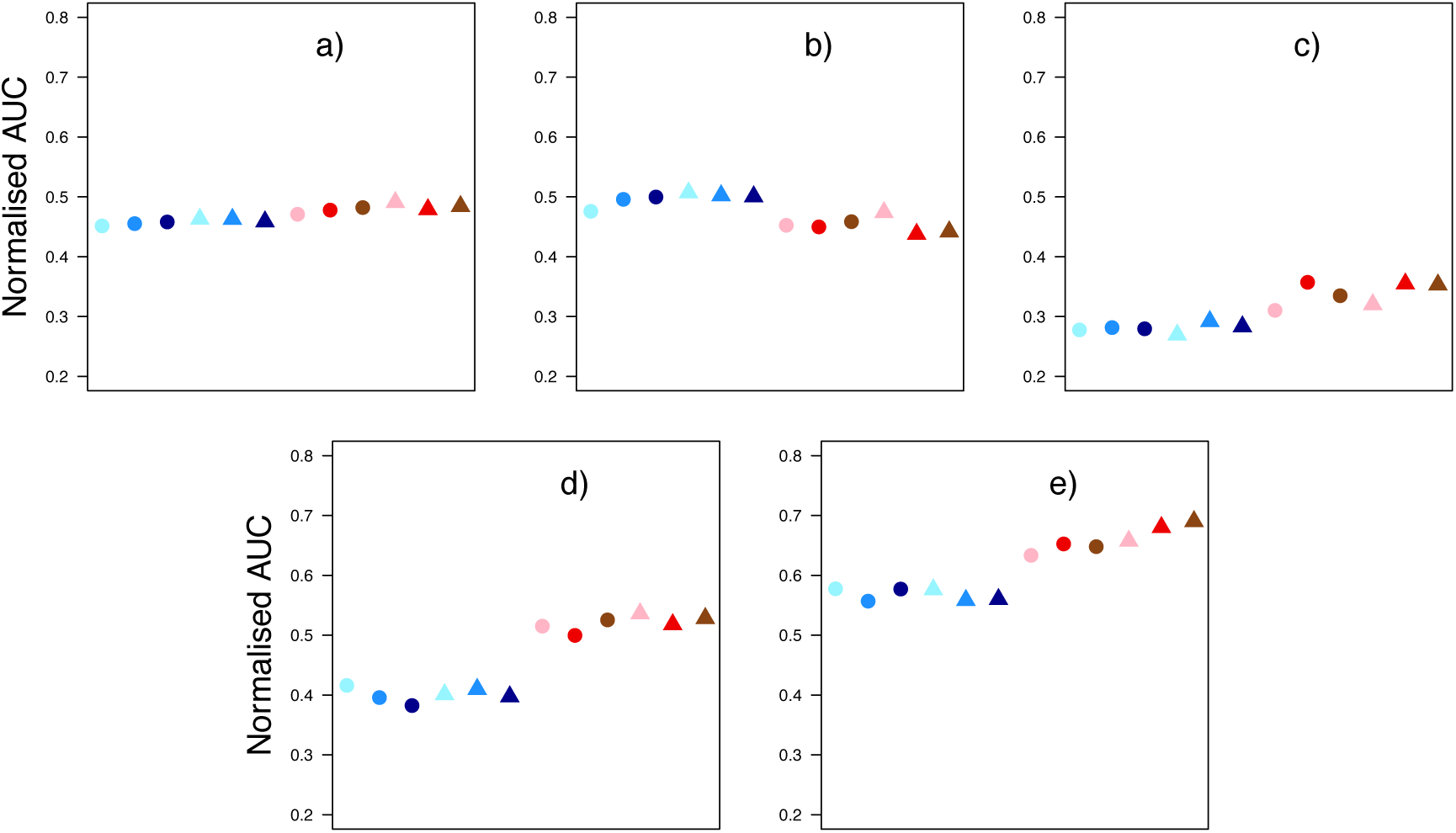
Area under curve (AUC, unitless) normalized by the duration of each period for each situation of interest, namely a combination of different levels of temperatures (blue: cold day; red: hot day), spatial disturbance (dot: low exposure; triangle: high exposure) and temporal disturbance (light colour: weekdays during school period; medium colour: weekdays during summer break; dark colour: weekends and public holidays): (a) total (1440 min), (b) diurnal (724.8 min), (c) nocturnal (235.2 min), (d) morning crepuscular (240 min) and (e) evening crepuscular (240 min) activity. AUC are calculated from predicted circadian activity patterns of northern chamois of the National Game and Hunting Reserve of Les Bauges massif, in summer (15 June to 7 September 2004–2020). We defined ‘diurnal’ as the period elapsed between 120 min after dawn and 120 min before dusk, ‘nocturnal’ as the period elapsed between 120 min after dusk and 120 min before dawn and ‘crepuscule’ as the remaining periods in the morning and in the evening (see text for details).

**Figure A8:**
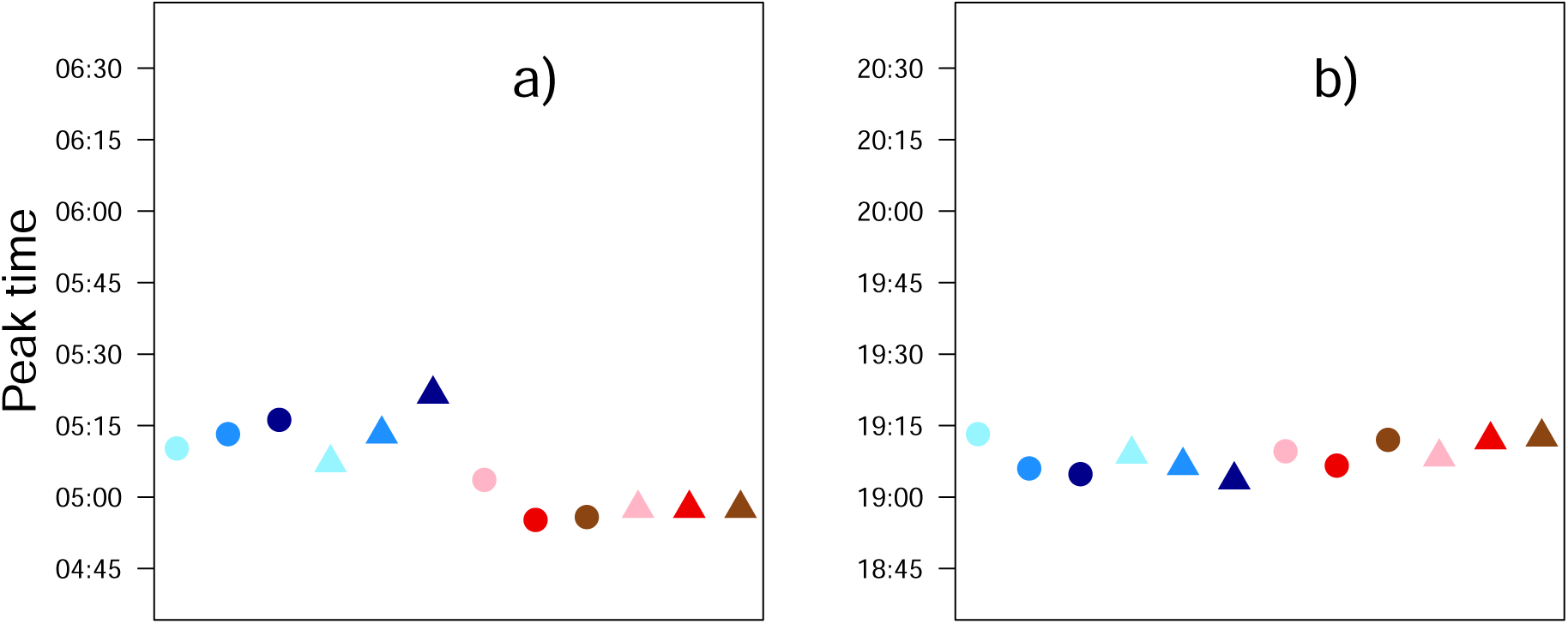
Timing of the activity peak (TAP, time of day when the activity peaks happen, i.e. the time when the highest probability of activity occurs) for each situation of interest, namely a combination of different levels of temperatures (blue: cold day; red: hot day), spatial disturbance (dot: low exposure; triangle: high exposure) and temporal disturbance (light colour: weekdays during school period; medium colour: weekdays during summer break; dark colour: weekends and public holidays): (a) morning peak and (b) evening peak. TAP are calculated from predicted circadian activity patterns of northern chamois of National Game and Hunting Reserve of Les Bauges massif, in summer (15 June to 7 September 2004–2020). Note the change of interval on the y axis between (a) and (b), although the scale stays the same.

**Figure A9:**
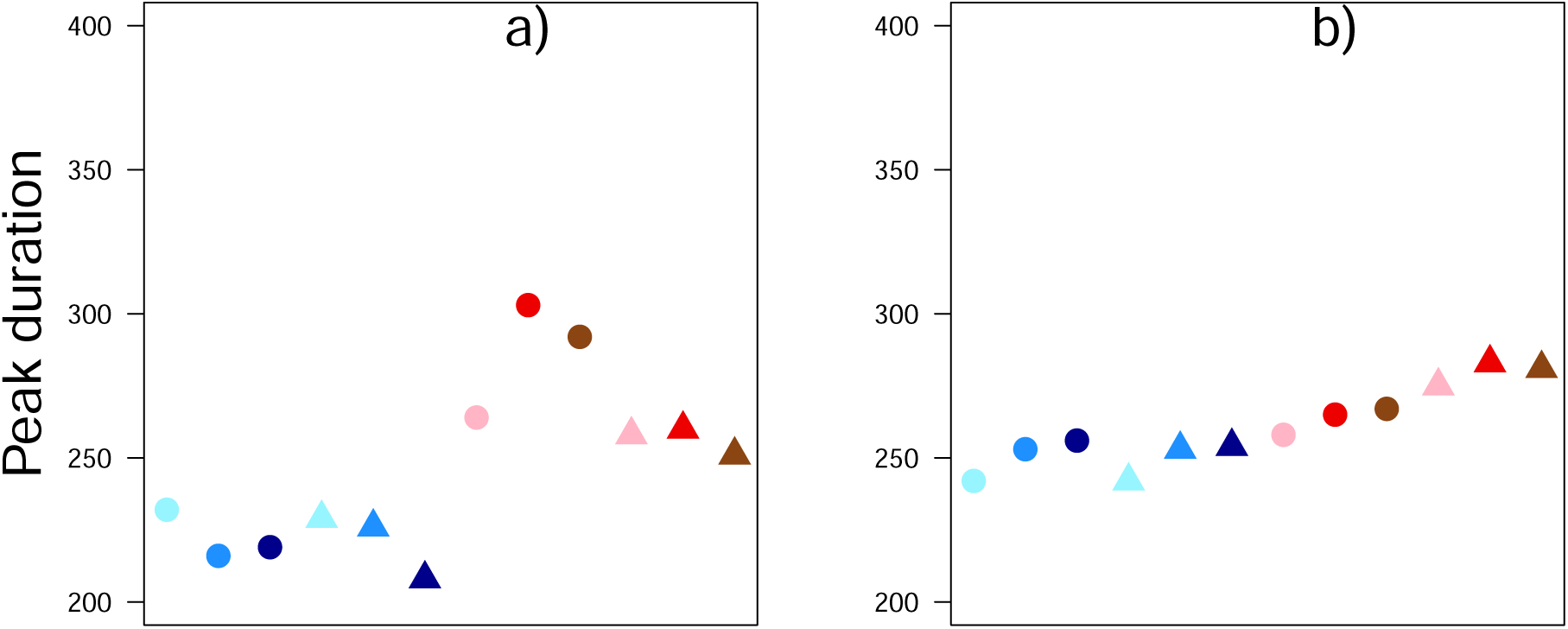
Peak duration (PD, min) for each situation of interest, namely a combination of different levels of temperatures (blue: cold day; red: hot day), spatial disturbance (dot: low exposure; triangle: high exposure) and temporal disturbance (light colour: weekdays during school period; medium colour: weekdays during summer break; dark colour: weekends and public holidays):(a) morning and (b) evening peak. PD are calculated from predicted circadian activity patterns of northern chamois of National Game and Hunting Reserve of Les Bauges massif, in summer (15 June to 7 September 2004–2020).

**Figure A10:**
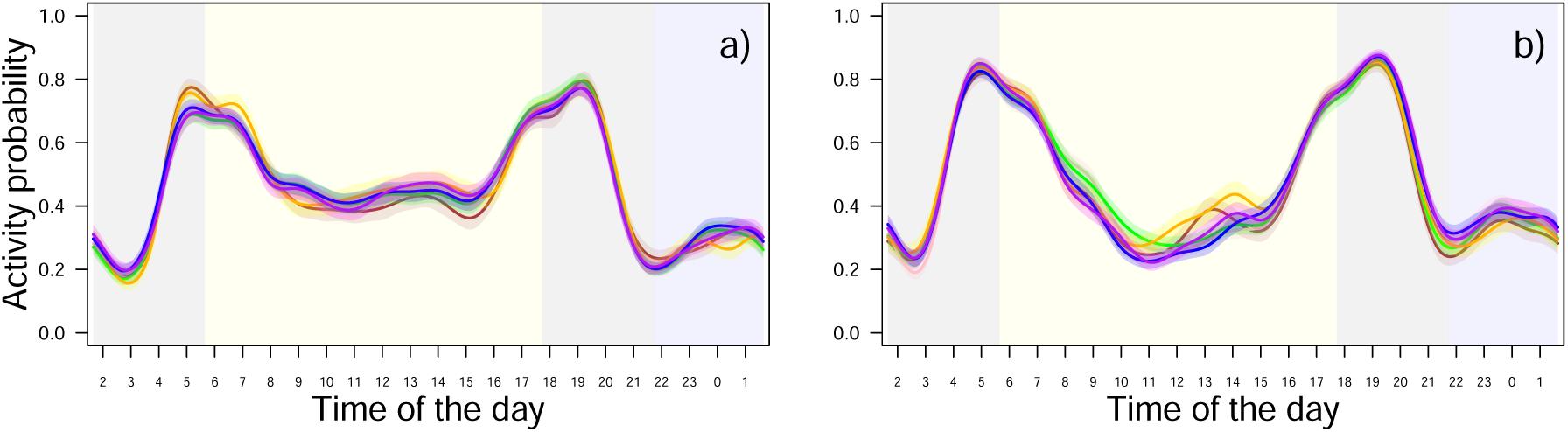
Predicted circadian activity patterns (activity ranges between 0 for inactivity and 1 for full-time activity) of female northern chamois in the National Game and Hunting Reserve of Les Bauges massif, in summer (15 June to 7 September 2004–2020), according to temperatures and human disturbance: (a) cold day and (b) hot day. In both (a) and (b), brown represents chamois living in the least exposed areas during weekdays of the school period; pink represents chamois living in the least exposed areas during weekdays of the summer break; green represents chamois living in the least exposed areas during weekends and public holidays; orange represents chamois living in the most exposed areas during weekdays of the school period; blue represents chamois living in the most exposed areas during weekdays of the summer break; purple represents chamois living in the most exposed areas during weekends and public holidays. Solid lines represent predicted values from the model and shaded areas around the curves represent 95% confidence intervals. Background shades represent the periods used to calculate the area under the curve for morning and evening crepuscules (grey), daytime (yellow) and night-time (blue). Time of the dayTime of day represents hours in Coordinated Universal Time (UTC).

**Figure A11:**
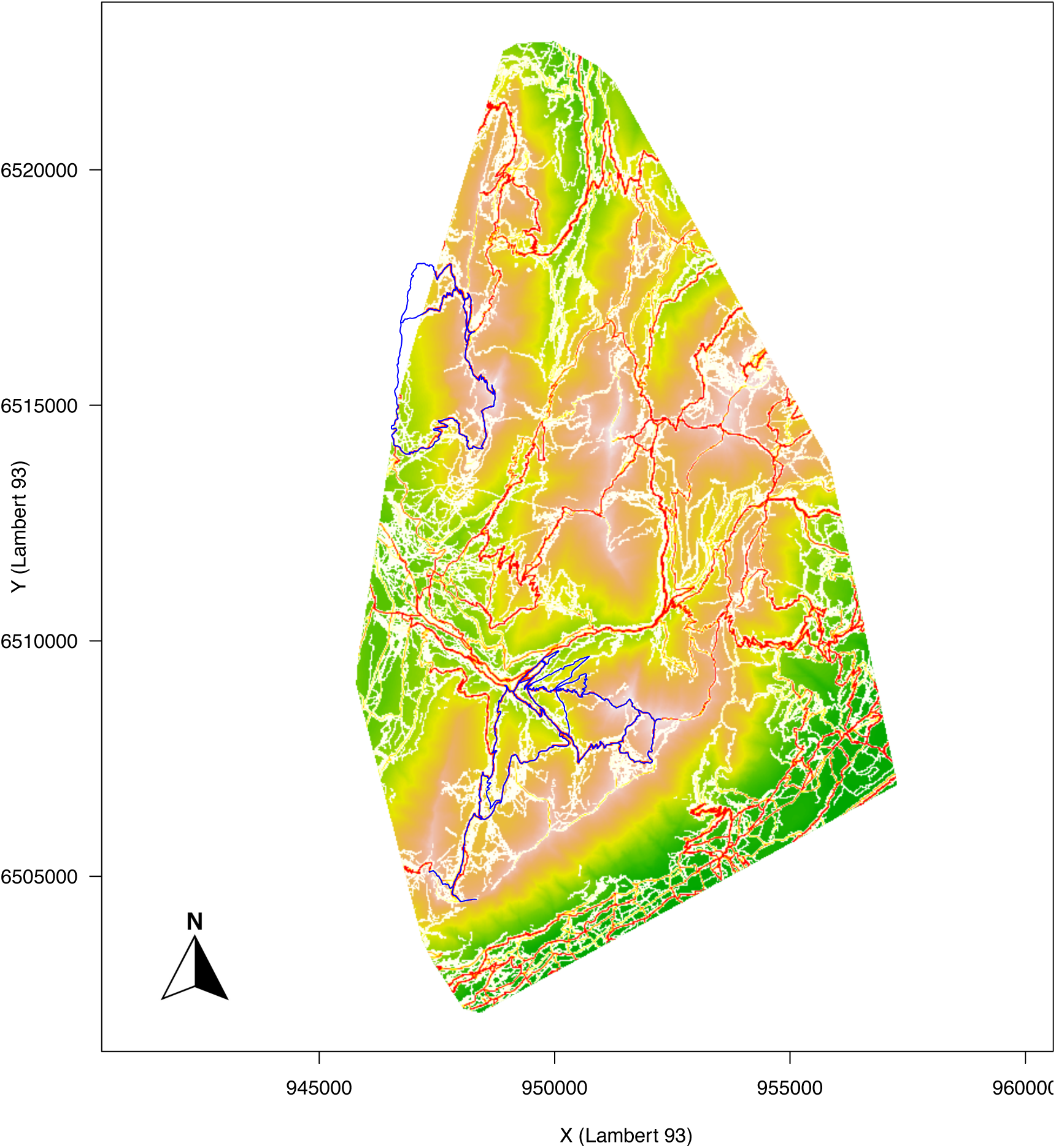
Comparison between human attendance assessed from hikers experimentally equipped with GPS in summer 2014 and 2015 (blue) and trail runners voluntarily uploading their GPS track on the Strava website (white-red gradient) in the National Game and Hunting Reserve of Les Bauges massif. Lambert 93: Lambert conformal conic projection, a conic map projection.

**Figure A12:**
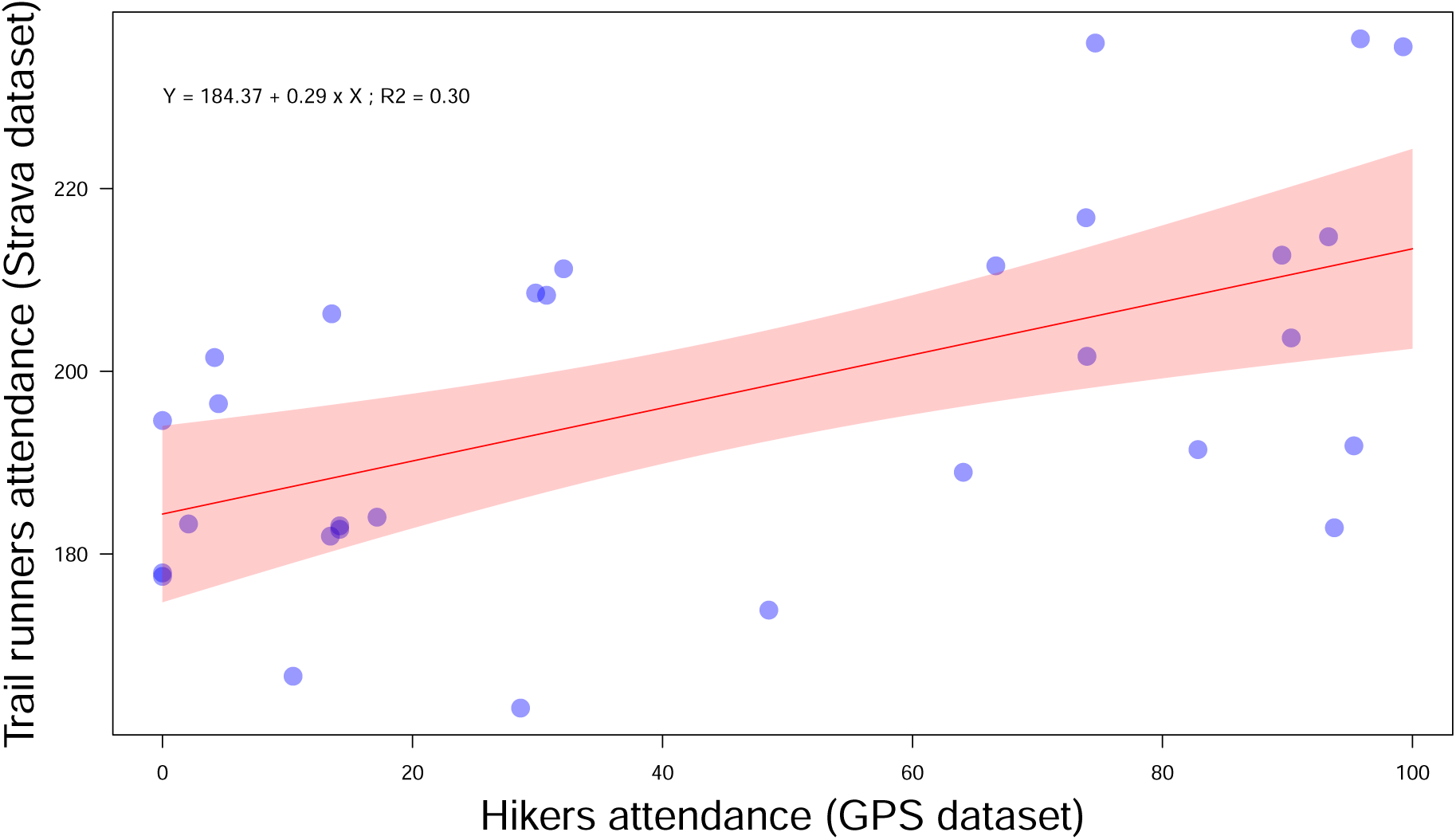
Comparison between human attendance assessed from hikers experimentally equipped with GPS trackers and from the Strava map in the National Game and Hunting Reserve of Les Bauges massif. Attendance levels assessed with the experimental design correspond to the proportion of use for each trail section, relative to each study site. Strava attendance levels correspond to the mean value of all the pixels of the Strava map located on a given trail segment of the map from the experimental design. The solid red line represents predicted values and the shaded area around it represents 95% confidence intervals.

**Figure A13:**
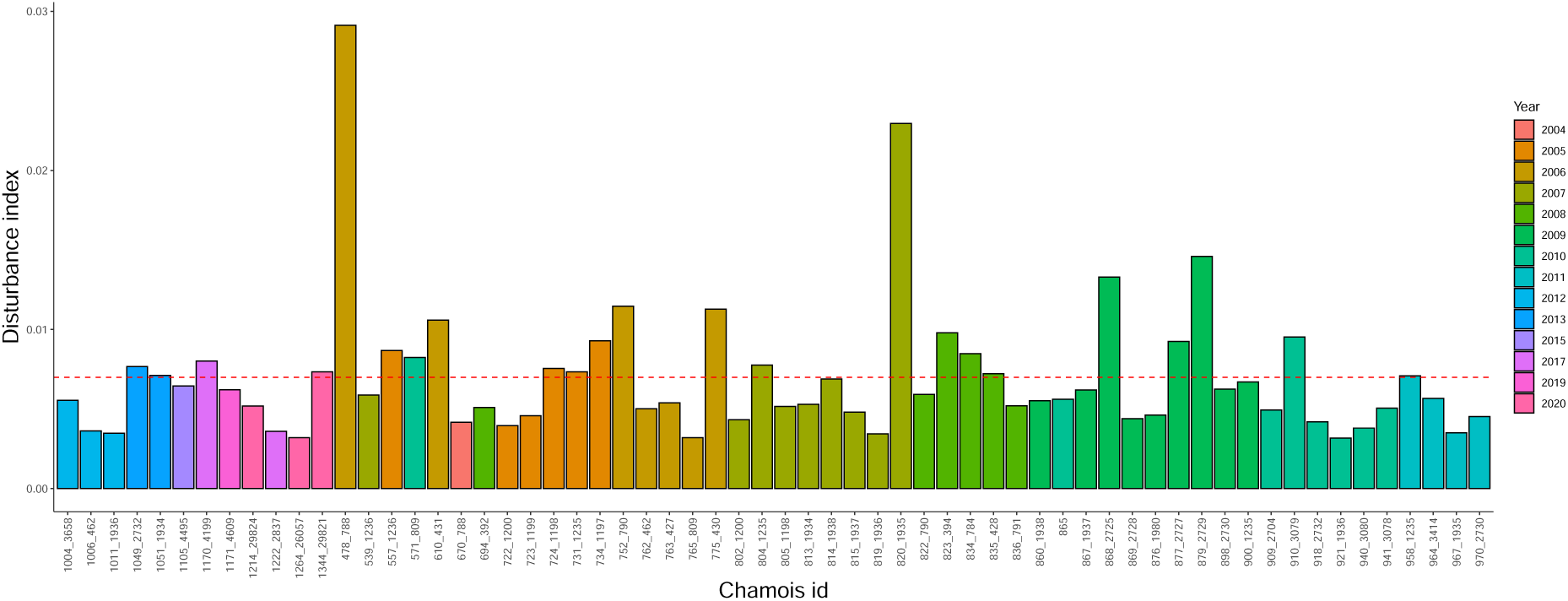
Spatial disturbance index per chamois home range for each year (the red horizontal dashed line corresponds to the median of the spatial disturbance index).

**Figure A14:**
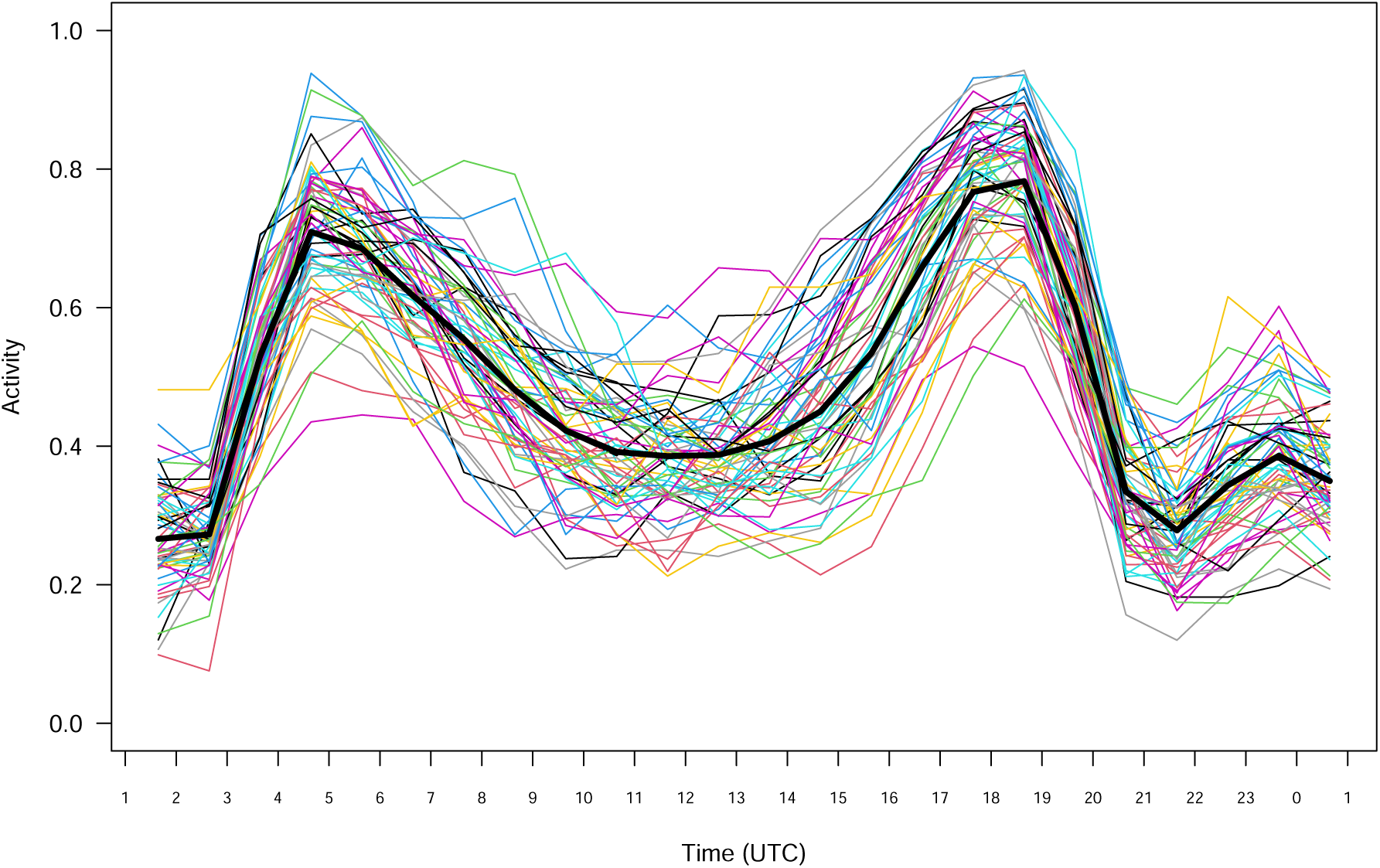
Mean hourly activity patterns (activity ranges between 0 for inactive and 1 for active) in summer for the northern chamois of the study (National Game and Hunting Reserve of Les Bauges massif, 15 June to 7 September 2004–2020). The mean activity probability for hour *H* was calculated from the 0/1 activity of the raw data ranging between *H* and *H+1* for a given chamois, considering all days available for this individual. Coloured solid lines represent the individual curves (*N* = 62). The thick black dotted line represents the general mean considering all the chamois and days available. Time is in hours, Coordinated Universal Time (UTC).

**Figure A15:**
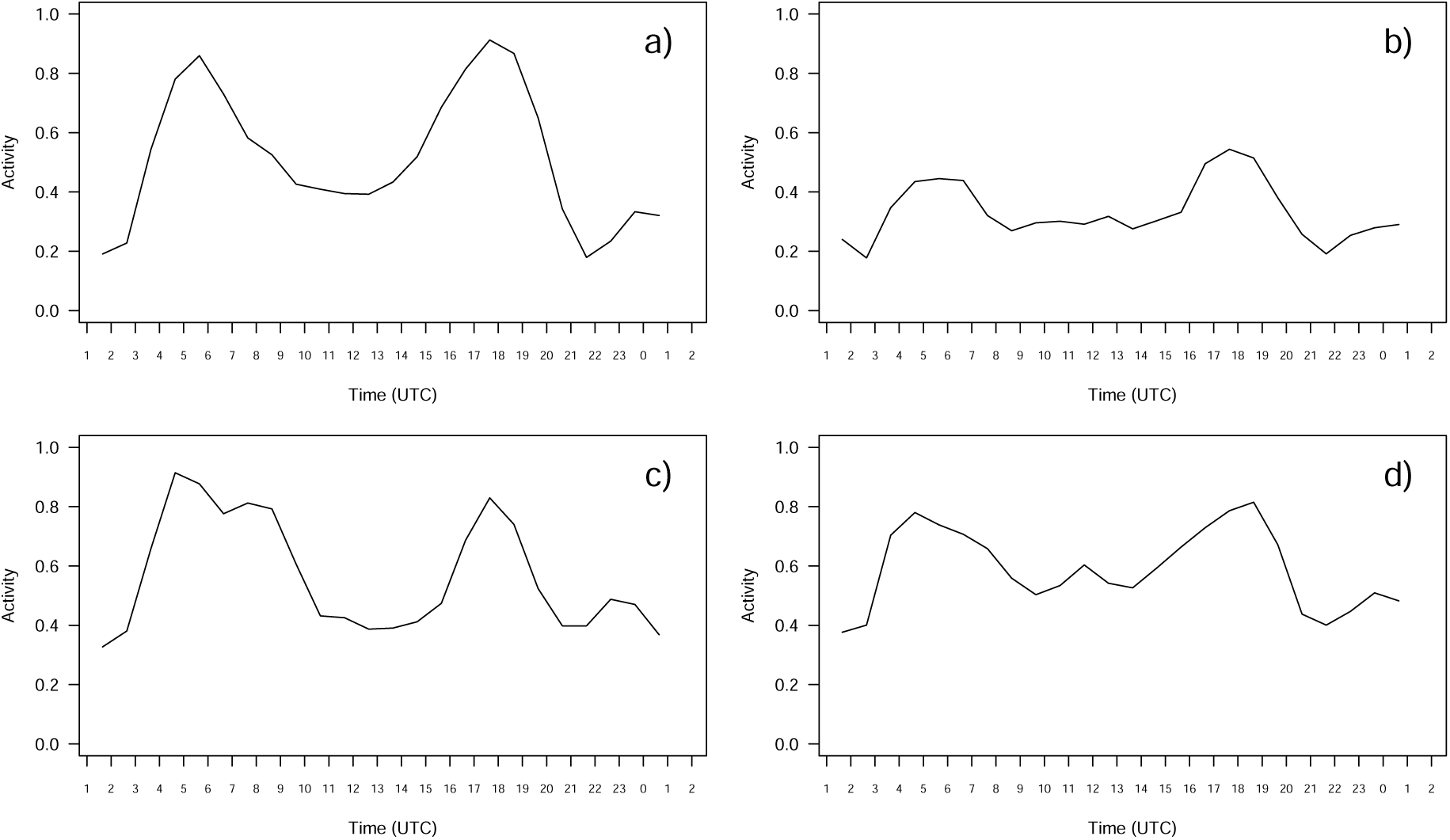
Mean hourly activity patterns (activity ranges between 0 for inactive and 1 for active) in summer for a selection of northern chamois from the National Game and Hunting Reserve of Les Bauges massif (15June to 7 September 2004–2020): (a) chamois 752_790 exhibits an activity pattern close to the mean pattern; (b) chamois 805_1198 exhibits an activity pattern characterized by small peaks; (c) chamois 1222_2837 exhibits an activity pattern characterized by a long morning peak; (d) chamois 834_784 exhibits an activity pattern characterized by a small additional peak during daytime. The mean activity probability for hour *H* was calculated from the 0/1 activity of the raw data ranging between *H* and *H+1*, considering all days available for this individual. Time is in hours, Coordinated Universal Time (UTC).

